# Bistable, biphasic regulation of PP2A-B55 accounts for the dynamics of mitotic substrate phosphorylation

**DOI:** 10.1101/2020.10.05.326793

**Authors:** Julia Kamenz, Lendert Gelens, James E. Ferrell

## Abstract

The phosphorylation of mitotic proteins is bistable, which contributes to the decisiveness of the transitions into and out of M phase. The bistability in substrate phosphorylation has been attributed to bistability in the activation of the cyclin-dependent kinase Cdk1. However, more recently it has been suggested that bistability also arises from positive feedback in the regulation of the Cdk1-counteracting phosphatase, PP2A-B55. Here, we demonstrate biochemically using Xenopus laevis egg extracts that the Cdk1-counteracting phosphatase PP2A-B55 functions as a bistable switch, even when the bistability of Cdk1 activation is suppressed. In addition, Cdk1 regulates PP2A-B55 in a biphasic manner; low concentrations of Cdk1 activate PP2A-B55 and high concentrations inactivate it. As a consequence of this incoherent feedforward regulation, PP2A-B55 activity rises concurrently with Cdk1 activity during interphase and suppresses substrate phosphorylation. PP2A-B55 activity is then sharply downregulated at the onset of mitosis. During mitotic exit Cdk1 activity initially falls with no obvious change in substrate phosphorylation; dephosphorylation then commences once PP2A-B55 spikes in activity. These findings suggest that changes in Cdk1 activity are permissive for mitotic entry and exit, but the changes in PP2A-B55 activity are the ultimate trigger.

## Introduction

The dramatic events of mitosis, including chromosome condensation, nuclear envelope breakdown, and spindle assembly, are driven by the collective phosphorylation of hundreds of proteins [1]. Dephosphorylation of these proteins promotes the transition out of mitosis. The majority of mitotic phosphorylations are mediated by cyclin-dependent kinase 1 (Cdk1) [2–4] and counteracted by a number of phosphatases, of which PP1 and PP2A-B55 have emerged as particularly important [5–8].

Until recently, the changes in substrate phosphorylation that drive mitotic entry and exit were mainly viewed in terms of the changing activity of Cdk1. Cdk1 gradually rises in activity during interphase due to cyclin synthesis [9–12]. Cdk1 activity then spikes at the onset of M phase as a result of the toggling of a bistable circuit [13–15] involving Wee1/Myt1, negative regulators of Cdk1 [16–20], and the Cdk1 activator Cdc25 [21, 22], from a high Wee1/low Cdc25 state to a low Wee1/high Cdc25 state. Once Cdk1 is fully activated, mitotic entry occurs. Shortly thereafter, the Cdc20-bound form of the anaphase-promoting complex/cyclosome (APC/C^Cdc20^) [23, 24] becomes activated and tags the mitotic cyclins for rapid degradation by the proteasome [25–27]. This inactivates Cdk1 and mitotic exit occurs.

However, it is now appreciated that along with these changes in kinase activity, there are marked changes in the activities of PP1 and PP2A-B55; both are negatively regulated by Cdk1 activity and therefore decrease in activity during M phase [6, 8, 28–31]. Moreover, the regulation of both phosphatases involves positive or double-negative feedback that could make them act as bistable switches. Indeed, based on computational modelling, Novak and colleagues proposed that at least PP2A-B55 functions as a bistable switch [32]. This hypothesis has been supported by in vitro reconstitution experiments. A system comprising Cdk1, PP2A-B55, and proteins that are part of the double-negative feedback loop regulating PP2A-B55 activity produces bistability in the phosphorylation of a model substrate even if Cdk1 activity does not exhibit bistability [33]. Furthermore, bistability in mitotic entry and exit was observed in human cells in response to the titration of Cdk1 activity. This bistability was only abolished when both the Cdk1and PP2A-B55 feedback loops were impaired [34].

In view of these findings, we have set out to determine the precise roles of kinase and phosphatase regulation in triggering the dramatic events of mitotic entry and exit. To this end we have used *Xenopus laevis* egg extracts, which can be made to proceed through the cell cycle *ex cellulo* and which are unusually amenable to biochemical analysis and manipulation. Our main findings are that (1) PP2A-B55 gradually increases in activity during interphase, suppressing the phosphorylation of mitotic phosphoproteins; (2) PP2A-B55 does, as hypothesized, function as a bistable switch, with its activity changing about 20-fold between its on state and its off state; (3) in quantitative terms, mitotic entry is driven primarily by phosphatase inactivation; and (4) mitotic exit is driven primarily by phosphatase reactivation. These findings suggest that changes in Cdk1 activity are permissive for mitotic entry and exit, but changes in phosphatase activity are instructive.

## Results

### Cdk1 substrate phosphorylation is bistable even when Cdk1 activity is not

Previous work in *Xenopus* extracts has shown that the activity of Cdk1 and the phosphorylation states of its substrates exhibit bistability and hysteresis [35, 36]. At low concentrations of cyclin B the system settles into a state of low Cdk1 activity and low substrate phosphorylation independently of the initial condition – the system is monostable. Similarly, high concentrations of cyclin B always force the system to settle into a state of high Cdk1 activity and high substrate phosphorylation – again the system is monostable. But at intermediate cyclin B concentrations, either of two alternative stable steady states is possible – the system is bistable. At these intermediate concentrations, the initial condition (the history of the system) determines which state the system is going to settle into: low Cdk1 activity/low phosphorylation if the system comes from interphase, or high Cdk1 activity/high phosphorylation if the system comes from M phase [35, 36]. It seemed reasonable to assume that the observed bistability in Cdk1 activity is causal to the observed bistability in substrate phosphorylation.

Yet, there is evidence suggesting that bistability in mitotic substrate phosphorylation may persist even when bistability in Cdk1 activity is eliminated. Wee1 activity is only essential for the first mitotic division in the model organism *Xenopus laevis* and the Wee1-mediated inhibitory phosphorylations on Cdk1 are sharply downregulated in the later cell cycles, implying that bistability in Cdk1 activity is dispensable for the later embryonic cell cycles [37–39]. Mitotic trigger waves, which in theory require positive feedback, continue to propagate through *Xenopus laevis* egg extracts when Wee1 is maximally inhibited [40]. In fission yeast, Wee1 – essential in the wild type strain – becomes dispensable in strains that are driven by the expression of a cyclin B-Cdk1 fusion protein [41, 42]. In addition, several Cdk1 substrates exhibit sharper phosphorylation thresholds during mitotic entry in extracts, where presumably all of the physiological regulators of their phosphorylation are present, than they do *in vitro* with purified Cdk1 and a constitutive phosphatase [43, 44]. The sharp phosphorylation thresholds persist even when Wee1 and Myt1 are inhibited [33, 43–45].

These findings, together with the reported potential of PP2A-B55 to exhibit bistability in vitro [33], suggested that an additional bistable circuit acting on one or more of the Cdk1-opposing phosphatases helps to keep the G2/M transition switch-like even when Cdk1 activity is forced to be graded and monostable.

To deconvolve these possibilities, we set out to study substrate phosphorylation in the absence of Cdk1 bistability. To eliminate bistability in Cdk1 activation, we abolished the double-negative feedback regulation of Cdk1 by blocking the Wee1/Myt1-mediated inhibitory phosphorylations of Cdk1 at Thr 14 and Tyr 15 [16–20] using the Wee1/Myt1 inhibitor PD0166825 [46] (Figure 1A). Since Cdc25 acts only on Thr/Tyr-phosphorylated Cdk1, the positive feedback loop is inhibited as well (Figure 1A). We began with titration experiments to determine the concentration of PD0166285 required to make Cdk1 activation become graded and monostable. The IC_50_ value for Wee1/Myt1 inactivation in extracts was found to be 1.4 to 3 *µ*M, similar to values reported previously [40] (Figures S1A–S1F). Based on these results, we used inhibitor concentrations between 5 *µ*M and 25 *µ*M whenever we aimed to eliminate the bistability of Cdk1 regulation.

**Figure 1.**
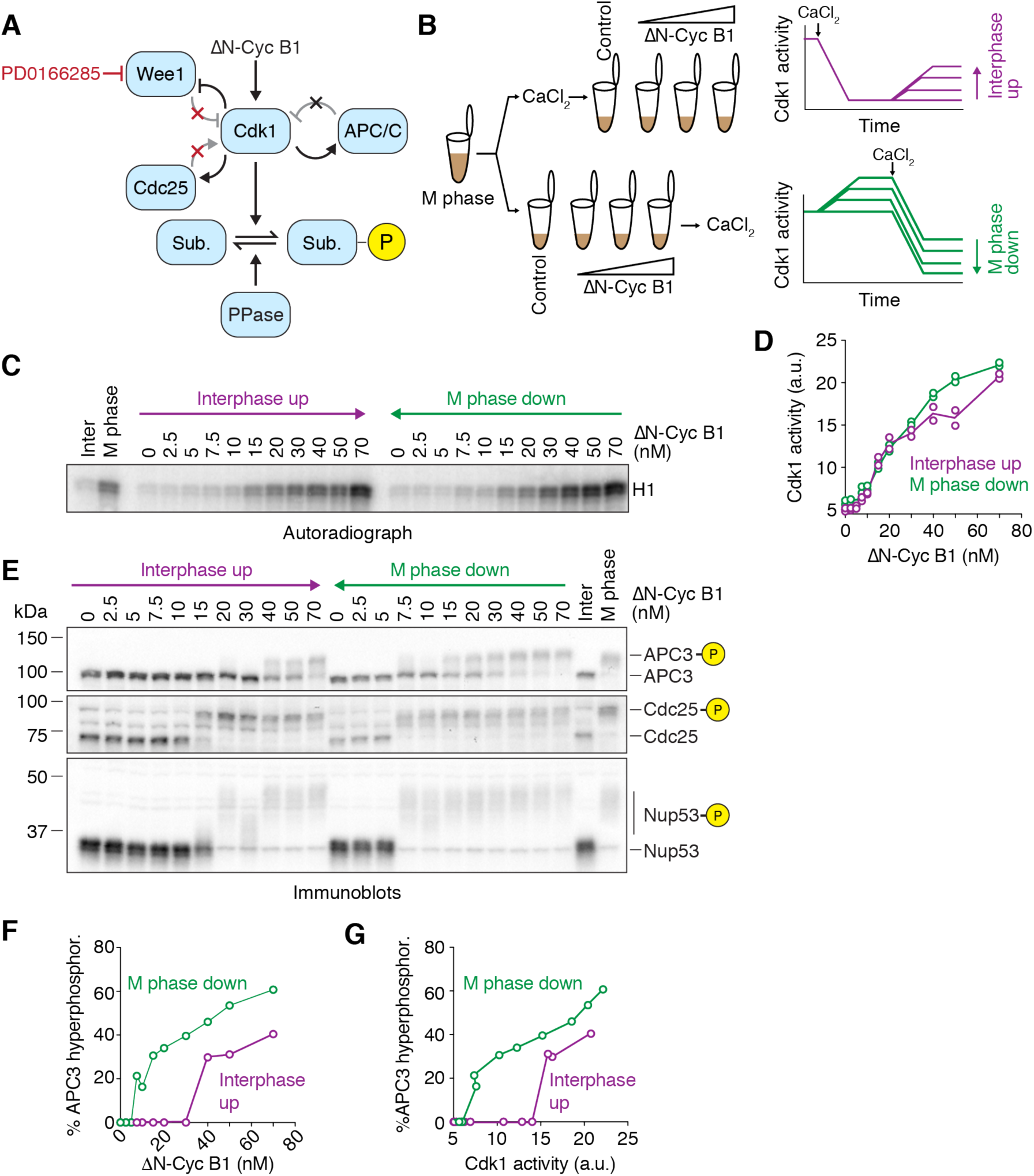
Cdk1 substrate phosphorylation is bistable in the absence of Cdk1 bistability. (A) Schematic of the regulation of substrate phosphorylation by Cdk1 and opposing phosphatases. Note that the Wee1 inhibitor PD0166285 compromises the positive and double negative feedback loops that regulate the activity of Cdk1. (B)Schematic of the hysteresis experiment. Steady-state Cdk1 activity and substrate phosphorylation were measured as a function of non-degradable cyclin B1 (ΔN-Cyc B1) in the presence of the Wee1/Myt1 inhibitor PD0166285. The steady state was approached either starting from a state of low (Interphase up, purple) or of high (M phase down, green) cyclin B concentration/Cdk1 activity. (C and D) Cdk1 activity as a function of ΔN-Cyc B1 concentration (as monitored by histone H1 phosphorylation) is graded and monostable in the presence of Wee1/Myt1 inhibitor (5 *µ*M PD0166285). Autoradiograph of the histone phosphorylation shown in (C). Quantification of two technical duplicates (circles) for the measurement and the mean of the duplicates (connecting lines) shown in (D). (E) The phosphorylation state of three Cdk1 substrates (APC3, Cdc25 and Nup53), monitored by the mobility shift as a function of ΔN-Cyc B1 concentration. Note that despite the monostable response in Cdk1 activity shown in (C) and (D) these substrates still exhibit hysteretic responses. (F-G) Quantification of the APC3 hyperphosphorylation for the experiment in (E), plotted as a function of ΔN-Cyc B1 concentration (F) and as a function of the corresponding Cdk1 activities (G) as measured in (D). Figure S1E shows a summary of multiple experiments for this measurement. All activity and phosphorylation state analyses (C)–(G) were performed from the same experiment. See Figure S1 for a detailed characterization of the Wee1 inhibitor (Figures S1A-S1F) and linearity tests for the antibodies (Figure S1H).

To look for hysteresis in substrate phosphorylation, we carried out experiments based on the general strategies that previously established the bistability of Cdk1 activation [35, 36]. We obtained M-phase arrested egg extracts (cytostatic factor (CSF)-arrested extracts), added cycloheximide to abolish cyclin synthesis and PD0166285 to inhibit Wee1 and Myt1, and treated the extracts in one of two ways. We supplemented one portion of the extract with 0.8 mM CaCl_2_ to induce the degradation of the endogenous cyclin B and reach low Cdk1 activity, and then added back different concentrations of non-degradable Δ65-cyclin B1. This way, the system approaches steady state from a point of low Cdk1 activity (Figure 1B, referred to as ‘interphase up’). Alternatively, we first added the Δ65-cyclin B1, which binds to excess Cdk1 in the extract and adopts the active phosphorylation state, and only afterward added CaCl_2_ to induce the selective degradation of the endogenous cyclin B. These extracts therefore approach steady state from initially high Cdk1 activity (Figure 1B, referred to as ‘M phase down’). After allowing the system to reach steady state, we measured Cdk1 activity and assessed the phosphorylation state of a set of well-characterized Cdk1 substrates. If the ‘interphase up’ and ‘M-phase down’ conditions yielded two different steady-state levels of substrate phosphorylation despite having a single steady-state level of Cdk1 activity, it would indicate that some phosphatase was bistable.

In the presence of the Wee1/Myt1 inhibitor, Cdk1 activity showed a graded, almost linear response to the titration of non-degradable cyclin B1 (Figures 1C and 1D). We consistently noted a small (∼5-10 nM) threshold in the response (Figures 1D and S1F). This may at least in part result from the presence of the Cdk inhibitor Kix1/Xic1, which has a reported concentration of ∼2 nM [47]. More importantly, the system now settled into a state of similar Cdk1 activity irrespective of the starting point, arguing that abolishing the positive and double-negative feedback eliminated the bistability of Cdk1 activity (Figures 1C and 1D), as expected.

We then asked whether hysteresis still persisted in the phosphorylation of three Cdk1 substrates (Figures 1E): APC3/Cdc27, a subunit of the anaphase-promoting complex/cyclosome (APC/C) considered a late substrate of Cdk1 during mitotic entry; the phosphatase Cdc25, an early substrate; and a subunit of the nuclear pore complex, Nup53/MP44, an intermediate (or middle) substrate [48]. All of these proteins exhibit phosphorylation-dependent changes in their electrophoretic mobility between interphase and M phase.

Although Cdk1 activity showed no hysteresis (Figures 1C and 1D), the three Cdk1 substrate proteins did (Figures 1E-G, and S1G). The cyclin B1 thresholds at which the substrates transitioned between hypo- and hyperphosphorylation differed from substrate to substrate, but in all cases the responses were hysteretic: higher cyclin B1 concentrations were needed to obtain hyperphosphorylation in the ‘Interphase up’ condition, than were needed to maintain hyperphosphorylation in the ‘M phase down’ condition. Therefore, substrate phosphorylation is hysteretic and bistable even when Cdk1 activity is not.

### APC/C activity is bistable in the absence of Cdk1 bistability

The Cdk1-mediated activation of APC/C^Cdc20^ is a crucial feature of the negative feedback at the core of the embryonic cell cycle oscillator [45, 49] (Figures 1A and 2A). An ultrasensitive or even bistable behavior in the activation of APC/C^Cdc20^ in response to Cdk1 activity could contribute to the robustness of this oscillator [45]. We had observed bistability in the hyperphosphorylation of the APC/C subunit APC3 (Figures 1E-G, and S1G), and this phosphorylation may be important for regulating the activity of the APC/C [49–51]. However, since the hyperphosphorylation of APC3 does not always correlate with the activity of the APC/C [45], we set out to directly measure the dose-response of APC/C activity as a function of the non-degradable cyclin B1 concentration.

**Figure 2.**
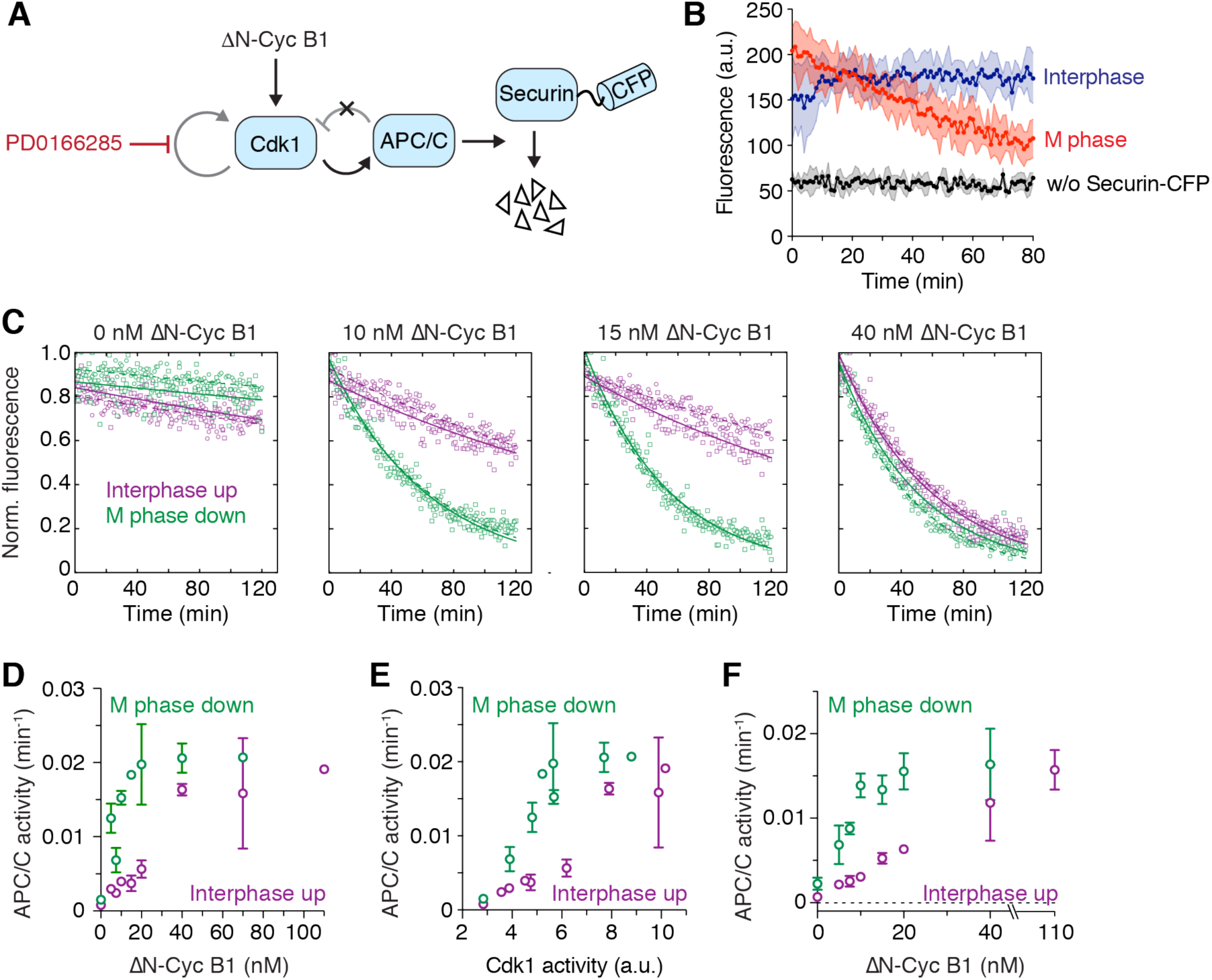
APC/C activity is bistable in the absence of Cdk1 bistability. (A) Schematic of the system used to assess APC/C activity. ΔN-Cyc B1 titration experiments were performed as described in Figure 1B and steady-state APC/C activity was measured by following the degradation of a fluorescently labeled APC/C substrate (securin-CFP) using a plate reader. (B) APC/C is activated during mitosis. Significant loss of fluorescent signal was only detected in an M phase extract (red), but not in an interphase extract (blue) or in an extract not supplemented with securin-CFP (black). Shown is the mean (circles with connecting line) and the standard deviation (error band) of a technical triplicate. (C) Securin-CFP degradation dynamics after adding different concentrations of ΔN-Cyc B1, approaching steady state starting from either a state of high (M phase down, green) or low (Interphase up, purple) cyclin B concentration/Cdk1 activity. Shown are data from two technical replicates (circles or squares, the dashed or solid lines, respectively, show the exponential fit of the data). Note that at intermediate concentrations of ΔN-Cyc B1 (10 or 15 nM), the steady-state level of APC/C activity depends upon whether the system has come from interphase or M phase. (D) Quantitation of the apparent first-order rate constant for APC/C activity plotted as a function of ΔN-Cyc B1 for the experiment shown in (C), including additional ΔN-Cyc B1 concentrations. Shown is the average of a technical duplicate with standard deviation. (E) Quantitation of the apparent first-order rate constant for APC/C activity (the same activities shown in panel (D)) plotted as a function of Cdk1 activity rather than non-degradable cyclin B concentration. Shown is the average of a technical duplicate with standard deviation. (F) Quantification of the apparent first-order rate constant for APC/C activity as a function of ΔN-Cyc B1. Shown are mean and standard error of the mean from 4 independent experiments (for 20 nM ΔN-Cyc B1: n=3, for 40 nM ΔN-Cyc B1: n=2). Note that due to variability between experiments the switch-like transition between low and high APC/C activity is less obvious in the averaged data than it is in the given single experiment (Figure 2D and 2E). All experiments were performed in the presence of 5 *µ*M PD0166285.

To this end, we performed a titration experiment similar to that described in Figure 1, but after allowing the extract to reach steady state, we added a fluorescently labeled substrate of the APC/C, securin-CFP, and followed substrate degradation using a fluorescence plate reader [45] (Figures 2A and 2B). The apparent rate constant for degradation serves as a gauge of APC/C activity.

As previously reported, in the absence of non-degradable Δ65-cyclin B1 (interphase), APC/C activity was low, whereas at high concentrations of Δ65-cyclin B1 (M phase), APC/C activity was high (Figure 2B). At low (0 nM) and high (40 nM) concentrations of Δ65-cyclin B1, the degradation kinetics did not depend on the initial starting conditions (Figures 2C-F). However, at intermediate concentrations of Δ65-cyclin B1 (e.g. 10 and 15 nM in Figures 2C and 2D), APC/C activity was high if the extract came from M phase and low if it came from interphase. This suggests that not only the phosphorylation state of APC/C but also the activity of APC/C is regulated in a hysteretic, bistable fashion even when Cdk1 activity is forced to be graded and monostable.

Note that protein synthesis is blocked in these experiments and therefore Emi2/XErp1, an additional regulator of APC/C activity which normally becomes synthesized during the first embryonic interphase [52] and could be a source for bistability in APC/C activity [53], is absent.

### PP1 does not exhibit bistability

Since substrate phosphorylation remained bistable even when Cdk1 activation was graded, we inferred it must result from bistability in the regulation of a counteracting phosphatase. Theoretically, Cdk1-mediated feedback regulation of either PP1 or PP2A-B55 could give rise to bistability (Figures 3A and 4A). In the case of PP1, two interlinked loops raise the possibility of bistability: Cdk1 inhibits PP1 by phosphorylating threonine 320 [28, 54, 55] and PP1 may activate itself by dephosphorylating T320 in an autocatalytic circuit. PP1 is further inhibited through binding to phosphorylated inhibitor-1 (I1) and can release itself from I1-mediated inhibition by dephosphorylation [8].

**Figure 3.**
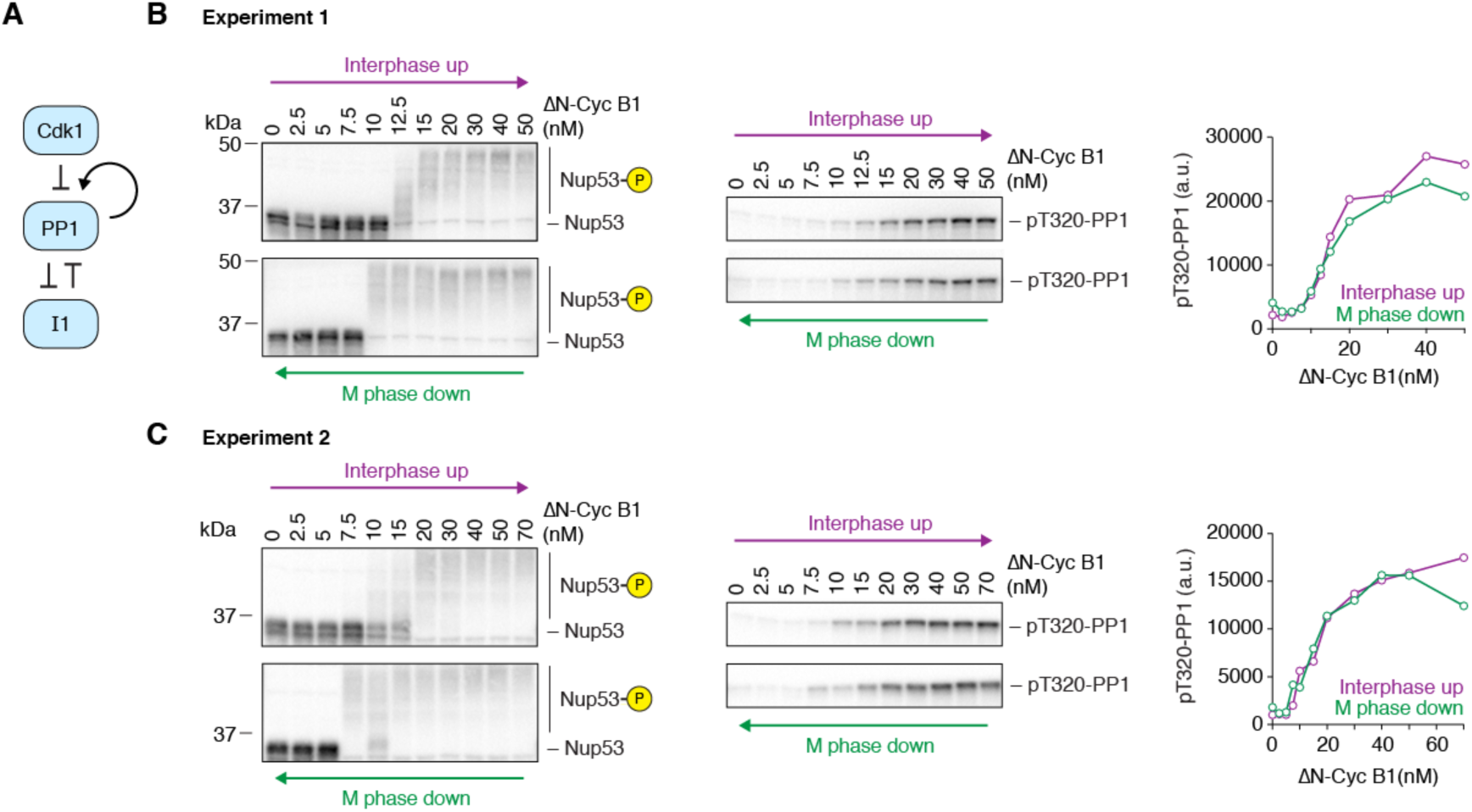
PP1 threonine 320 phosphorylation does not exhibit bistability. (A) Schematic of the regulation of PP1 activity by Cdk1 and inhibitor-1 (I1). (B and C) Two independent experiments examining whether the phosphorylation of the C-terminal regulatory tail of PP1 at threonine 320, which can be taken as a measure of PP1 activity, is hysteretic. Although Nup53 phosphorylation showed bistable behavior in both experiments, PP1 threonine 320 phosphorylation was graded and monostable. 10 *µ*M PD0166285 was used to inhibit Wee1/Myt1 in (B) and 5 *µ*M in (C).

**Figure 4.**
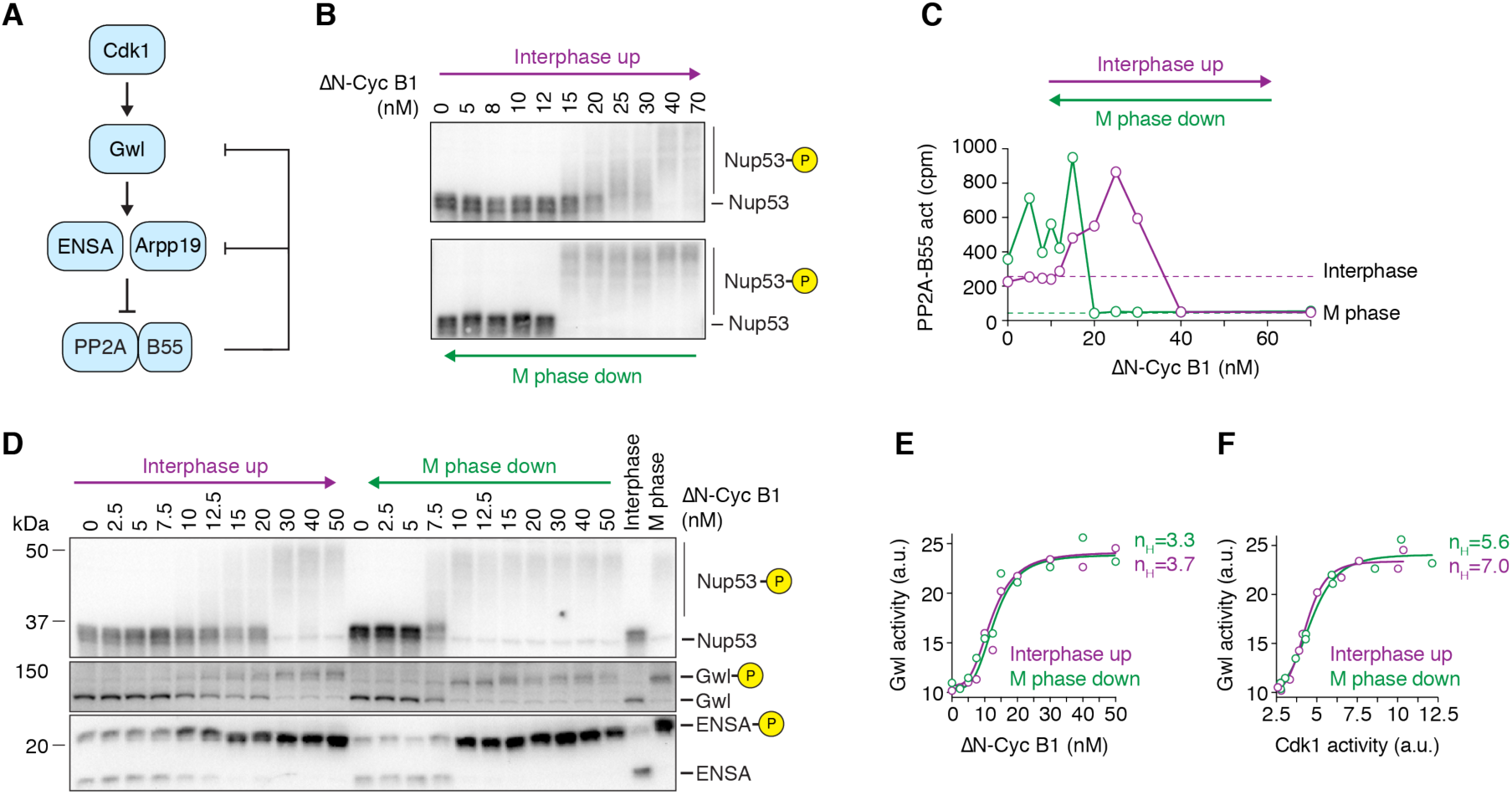
PP2A-B55 activity, but not Gwl activity, exhibits bistability when Cdk1 activity is graded and monostable. (A) Schematic of the regulation of PP2A-B55 activity by Cdk1 via Greatwall kinase (Gwl) and ENSA/Arpp19. Two double-negative feedback loops, a shorter one involving only PP2A-B55 and Arpp19/ENSA and a longer one involving Gwl kinase and PP2A-B55, could give rise to bistability. (B and C) The steady-state activity of PP2A-B55 is hysteretic in the presence of Wee1/Myt1 inhibitor (10 *µ*M PD0166285). Mitotic substrate phosphorylation (monitored by the mobility shift of Nup53) in (B) and PP2A-B55 activity in (C) are shown as a function of non-degradable cyclin B1 (ΔN-Cyc B1), approaching steady state from either a state of high (M phase down, green) or low (Interphase up, purple) cyclin B concentration/Cdk1 activity (mean of technical duplicates with connecting line). Two additional experiments are shown in Figures S2B-E. (D) Gwl and ENSA phosphorylation exhibit bistability in the presence of Wee1/Myt1 inhibitor (5 *µ*M PD0166285). Phosphorylation states of several Cdk1 substrates including Gwl and ENSA were analysed by immunoblotting. Figure S3A shows an additional experiment. 10 *µ*M Phos-tag was used to enhance the mobility shift upon ENSA phosphorylation. A shorter exposure for the ENSA immunoblot and a quantification of the phosphorylated form of ENSA are shown in Figures S3A and S3B. An additional experiment is shown in Figures S3C and S3D and an experiment detecting ENSA phosphorylation using a phospho-specific antibody against the phosphorylated S67 epitope of ENSA is shown in Figures S3I and S3J. (E and F) Greatwall kinase activity is ultrasensitive but not bistable. Cdk1 and Gwl kinase activities were measured for the experiment shown in (D) and depicted as the dose-response of Gwl kinase activity as a function of ΔN-Cyc B1 (E) or Cdk1 activity (F). Shown is the mean of a technical duplicate (circles). While not bistable, the dose-response of Greatwall kinase activity exhibits significant ultrasensitivity as demonstrated by the large Hill exponent (n_H_) necessary to fit the data (solid lines in (E) and (F)). Additional experiments are shown in Figures S3G-L.

Thus we set out to determine whether the response of PP1 to Cdk1 was bistable or monostable by using the phosphorylation state of threonine 320 as a proxy for PP1 activity; the higher the T320 phosphorylation, the lower the activity. In contrast to the Cdk1 substrates shown in Figures 1 and 2, PP1 T320 phosphorylation showed a linear, monostable response to Δ65-cyclin B1 concentration and Cdk1 activity, with little if any hysteresis (Figures 3B and 3C). This finding suggests that PP1 is not the phosphatase responsible for the observed bistability in substrate phosphorylation and APC/C activity (Figures 1 and 2). The lack of hysteresis in PP1 phosphorylation underscores the fact that the presence of positive feedback and double-negative feedback does not guarantee that a system will exhibit bistability.

### PP2A-B55 is regulated in a bistable, biphasic fashion

We next investigated whether PP2A-B55 exhibits bistability (Figure 4A) as hypothesized [32–34]. In mitosis Cdk1 phosphorylates and activates Greatwall (Gwl) [30, 56–58]. Active Gwl then phosphorylates two closely-related proteins, Arpp19 and ENSA [59, 60], which in their phosphorylated state bind and inhibit PP2A-B55 by an ‘unfair competition’ mechanism [61]. PP2A-B55 promotes its own release from this inhibition by dephosphorylating Arpp19/ENSA and by dephosphorylating Gwl [62–64], though PP1 also significantly contributes to Gwl dephosphorylation [63–65]. Therefore, there are at least two double negative feedback loops that regulate PP2A-B55 and could potentially generate bistability: one between PP2A-B55 and Greatwall, and one between PP2A-B55 and the stoichiometric inhibitors ENSA and Arpp19 (Figure 4A).

We measured PP2A-B55 activity based on the release of inorganic phosphate from a ^32^P-labeled peptide encompassing the serine 50 site of Cdc20 [66] (Figure S2A). The specificity of this assay has been established through immunodepletion experiments, which showed that PP2A-B55, and in particular PP2A-B55*δ*, accounts for most of the phosphatase activity toward this substrate in *Xenopus* extracts [6]. As previously reported [6], PP2A-B55 activity was significantly higher in interphase than in M phase (Figure S2A).

We then carried out a hysteresis experiment and measured mitotic substrate phosphorylation (Figure 4B) as well as PP2A-B55 activity (Figure 4C). In the ‘Interphase up’ leg of the experiment, PP2A-B55 activity was found to increase about 4-fold as the Δ65-cyclin B1 was raised from 0 to 25 nM, and it then decreased about 20-fold once mitotic concentrations (40 nM) of cyclin were reached (Figure 4C). Thus the response of PP2A-B55 to Δ65-cyclin B1 was biphasic, with low Cdk1 activities positively regulating the phosphatase and high Cdk1 activities negatively regulating it.

The response was also hysteretic, with a higher concentration of Δ65-cyclin B1 required to inhibit PP2A-B55 coming from interphase than was required to maintain its inactivation coming from M phase (Figure 4C, purple vs. green curves). Note that the biphasic response was seen in both the ‘Interphase up’ and ‘M phase down’ legs of the experiment (Figure 4C), and that both the biphasic response and the bistability were consistently observed (Figures S2B–S2E).

To determine whether the double-negative feedback between PP2A-B55 and Gwl and/or PP2A-B55 and ENSA/Arpp19 was important for the observed bistability in PP2A-B55 activity, we assessed the phosphorylation state of Gwl and ENSA/Arpp19. As shown in Figures 4D, S3C and S3I, the hyperphosphorylation of Gwl was hysteretic under conditions where PD0166285 had eliminated hysteresis in Cdk1 activity, suggesting that the PP2A-B55-Gwl double-negative feedback might be critical for the bistability of PP2A-B55. Similarly, we detected some hysteresis in the phosphorylation response of ENSA and Arpp19, either by resolving the different phosphorylation forms using Phos-tag gels (Figure 4D and S3A-D) or using a phospho-specific antibody (Figure S3I and S3J) raised against the Gwl target sites serine 67 and serine 62 in ENSA and Arpp19, respectively.

These findings suggested that the PP2A-B55-Gwl double-negative feedback might be critical for the bistability of PP2A-B55. However, although PP2A-B55 has been shown to dephosphorylate Gwl, it has recently been shown that the activity of Gwl is mainly regulated by PP1 rather than PP2A-B55 [63–65]. Furthermore, the hyperphosphorylation of Gwl only weakly correlates with its activity [63]. Therefore, we asked whether the observed bistability in Gwl phosphorylation actually translated into bistability in Gwl activity.

To measure Gwl activity, we used a recombinant Arpp19 protein that carried alanine mutations in two prominent non-Gwl phosphorylation sites (S28 and S109, Arpp19-2A) as an *in vitro* substrate and monitored incorporation of radioactive phosphate from [*γ*-^32^P]ATP, as previously described ([63], Figures S3E and S3F). Although we had clearly detected hysteresis in the phosphorylation state of Gwl, we did not detect hysteresis in Gwl activity. The response curves were ultrasensitive, but indistinguishable for extracts coming out of interphase versus M phase (Figures 4E, 4F, S3G, and S3H). Note that in our enzymatic assays to monitor Gwl activity we used okadaic acid, a PP2A inhibitor. Although this is commonly done in order to preserve the phosphorylation and activity state of Gwl during the measurement, it could have altered the outcome of the measurement. However, even when we isolated Gwl from extracts and measured Gwl activity in the absence of okadaic acid, we did not detect bistability in Gwl activity (Figure S3I-L).

We cannot formally exclude the possibility that our assay is too noisy to detect some small degree of bistability in Gwl activity. However, given that we did not detect bistability in PP1 activity and PP1 is thought to be an important regulator of Gwl activity, we favor the possibility that bistability in ENSA/Arpp19 phosphorylation and PP2A-B55 activity does not arise from double-negative feedback between PP2A-B55 and Gwl, but that instead the feedback interaction between PP2A-B55 and Arpp19/ENSA is responsible for the observed hysteresis in PP2A-B55 regulation.

### Partially inhibiting PP2A activity abolishes the bistability in Cdk1 substrate phosphorylation

If indeed the bistability of PP2A-B55 activity is responsible for the observed bistability in substrate phosphorylation and APC/C activity, then partially inhibiting PP2A-B55 might abolish this bistable behavior. This could either be a result of disturbing the balance between the two legs of the PP2A-B55-Arpp19/ENSA double-negative feedback loop or because phosphatases other than PP2A become dominant in regulating the phosphorylation of these substrates when PP2A-B55 is inhibited. We first chose to decrease PP2A-B55 activity using okadaic acid (OA) (Figure 5A). OA is a high affinity inhibitor of PP2A with a half maximal inhibitor concentration (IC_50_) of about 0.14 nM in vitro [67] but also inhibits PP4 and at higher concentrations PP1 and PP5 [60]. Therefore, we carried out titration experiments to identify a concentration of OA where the balance between Cdk1-mediated phosphorylation and PP2A-B55-mediated dephosphorylation was slightly shifted towards the phosphorylation reaction, but PP1 activity was not impaired. Concentrations of 0.4 *µ*M to 0.6 *µ*M OA induced subtle changes in the phosphorylation states of several Cdk1 substrates (Cdc25, Nup53, Wee1, and Gwl, but not APC3) in the presence of low amounts of non-degradable cyclin B1, but did not affect the phosphorylation state of T320 of PP1 (Figure S4A), suggesting that at these concentrations we had partially inhibited PP2A but not PP1. Accordingly, we set out to see whether these low concentrations of OA would have any effect on the hysteresis seen in Cdk1 substrate phosphorylation.

**Figure 5.**
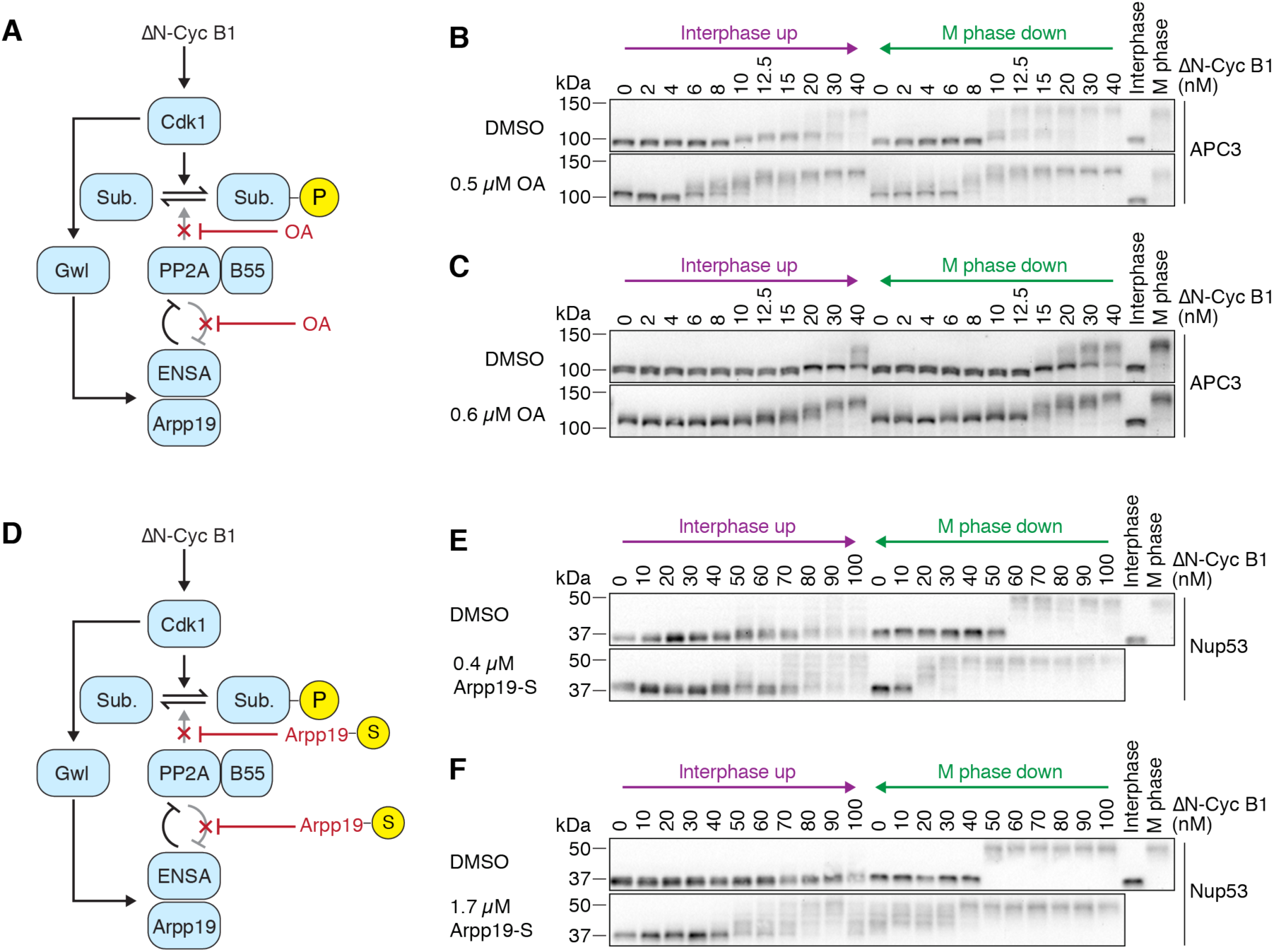
Dampened PP2A activity weakens the bistability in Cdk1 substrate phosphorylation. (A) Okadaic acid (OA) was used to partially inhibit the activity of PP2A-B55 and the feedback between PP2A-B55 and ENSA/Arpp19. (B and C) The dose-response relationship of the phosphorylation state of the APC/C subunit APC3 as a function of ΔN-Cyc B1, approaching steady-state starting from either a state of high (M phase down) or low (Interphase up) cyclin B concentration/Cdk1 activity in the presence of the Wee1/Myt1 inhibitor (25 *µ*M PD0166285) and in the absence (DMSO) or presence (OA) of different concentrations of OA. OA decreased the hysteresis seen in the DMSO control and made intermediate phosphorylation states more apparent. Note that the DMSO and OA comparison were performed in parallel using the same extract, but (B) and (C) were performed using different extracts; biological variability may account for the differences in the overall response between the two DMSO control experiments. (D) Thio-phosphorylated Arpp19 (Arpp19-S) was used to partially inhibit the activity of PP2A-B55 and the feedback between PP2A-B55 and ENSA/Arpp19. (E and F) The dose-response relationship of the phosphorylation state of Nup53 as a function of ΔN-Cyc B1, approaching steady-state starting from either a state of high (M phase down) or low (Interphase up) cyclin B concentration/Cdk1 activity in the presence of the Wee1/Myt1 inhibitor (10 *µ*M PD0166285) and in the absence (DMSO) or presence (Arpp19-S) of different concentrations of Arpp19-S. Arpp19-S decreased the hysteresis seen in the DMSO control and made intermediate phosphorylation states more apparent. Note that the DMSO and Arpp19-S comparison were performed in parallel using the same extract, but and (F) were performed using different extracts; biological variability may account for the differences in the overall response between the two DMSO control experiments, and variability in the activity of the ΔN-Cyc B1 preparation might account for overall differences in the response curve when comparing C and D to E and F.

In the presence of the Wee1 inhibitor PD0166285, 0.5 *µ*M OA changed the dose-response of APC3 phosphorylation in three ways. First, the dose-response curve shifted towards lower non-degradable cyclin B1 concentrations; on the ‘Interphase up’ leg half-maximal phosphorylation was obtained at a cyclin concentration of ∼8 nM, compared with 30-40 nM in the absence of OA (Figure 5B, compare top versus bottom panel). Second, the phosphorylation of APC3 became more graded; whereas in the absence of OA APC3 exhibited mainly two prominent phosphorylation states (a hypo- and a hyperphosphorylated state), addition of OA yielded a variety of APC3 forms of intermediate mobility in a dose-dependent manner (Figure 5B). Finally, the bistability in the dose-response relationship was almost completely abolished (Figure 5B); the phosphorylation responses starting in interphase and starting in M phase were very similar. Increasing the OA concentration to 0.6 *µ*M potentiated all of these effects (Figure 5C).

The thiophosphorylated form of Arpp19 has been shown to be a specific, high-affinity inhibitor of PP2A-B55 with an IC_50_ of 0.47 nM in vitro [61, 68]. We therefore used thiophosphorylated Arpp19 (Arpp19-S) as an alternative way of inhibiting PP2A-B55 and perturbing the feedback regulation (Figure 5D). We used nominal concentrations between 0.4 *µ*M and 1.7 *µ*M of Arpp19-S to inhibit PP2A-B55 (though the actual inhibitor concentration was estimated to be about 50% lower due to incomplete thiophosphorylation of Arpp19). In the presence of 25 nM non-degradable cyclin B1, addition of Arpp19-S but not unphosphorylated Arpp19 induced significant changes in the phosphorylation state of Nup53 and Gwl (Figure S4B). These concentrations are consistent with stochiometric inhibition of PP2A-B55 in the extract, where the catalytic subunit of PP2A is present at about 0.3 *µ*M [69]. At higher concentrations of non-degradable cyclin B1, both thiophosphorylated and unphosphorylated Arpp19 were able to induce changes in the phosphorylation states of Nup53 and Gwl with the thiophosphorylated Arpp19 being more potent. Similar results were reported previously [68].

When comparing the dose-response of substrate phosphorylation with or without Arpp19-S in the presence of the Wee1 inhibitor PD0166285, we observed effects similar to the ones we had observed with low doses of OA. In particular, we observed that in the presence of Arpp19-S, half-maximal substrate phosphorylation occurred at lower concentrations of cyclin B1, and that the response overall seemed more graded with more intermediate phosphorylation bands being detected (Figures 5E and 5F). The impact of Arpp19-S on bistability was less obvious in these experiments than it had been using OA. These findings leave open the possibility that some phosphatase in addition to PP2A-B55 contributes to the observed hysteresis.

### Both the Cdk1 and the PP2A switch contribute to the bistability in substrate phosphorylation

So far we have shown that PP2A-B55 regulation is bistable even when Cdk1 activity is not bistable. However, in the absence of PD0166285, and in the first cell cycle of an embryo, both the Cdk1 switch and the PP2A-B55 switch should be operative. The relative contributions of the two switches to substrate phosphorylation are unclear; therefore, we asked whether any hysteresis would remain when the PP2A-B55 loop was compromised by OA treatment but the Cdk1/Wee1/Cdc25 system was not compromised by PD0166285 treatment.

To this end, we attenuated PP2A activity using 0.5 *µ*M OA and compared the dose-response of substrate phosphorylation in the presence or absence of Wee1 inhibition. In the presence of OA (but no PD0166285), substrate phosphorylation was hysteretic (Figures 6A and 6B), whereas OA plus PD0166285 abolished hysteresis (Figures 6C and 6D) as we had observed previously (Figure 5B). We conclude that both the kinase switch and the phosphatase switch contribute to bistability of substrate phosphorylation.

**Figure 6.**
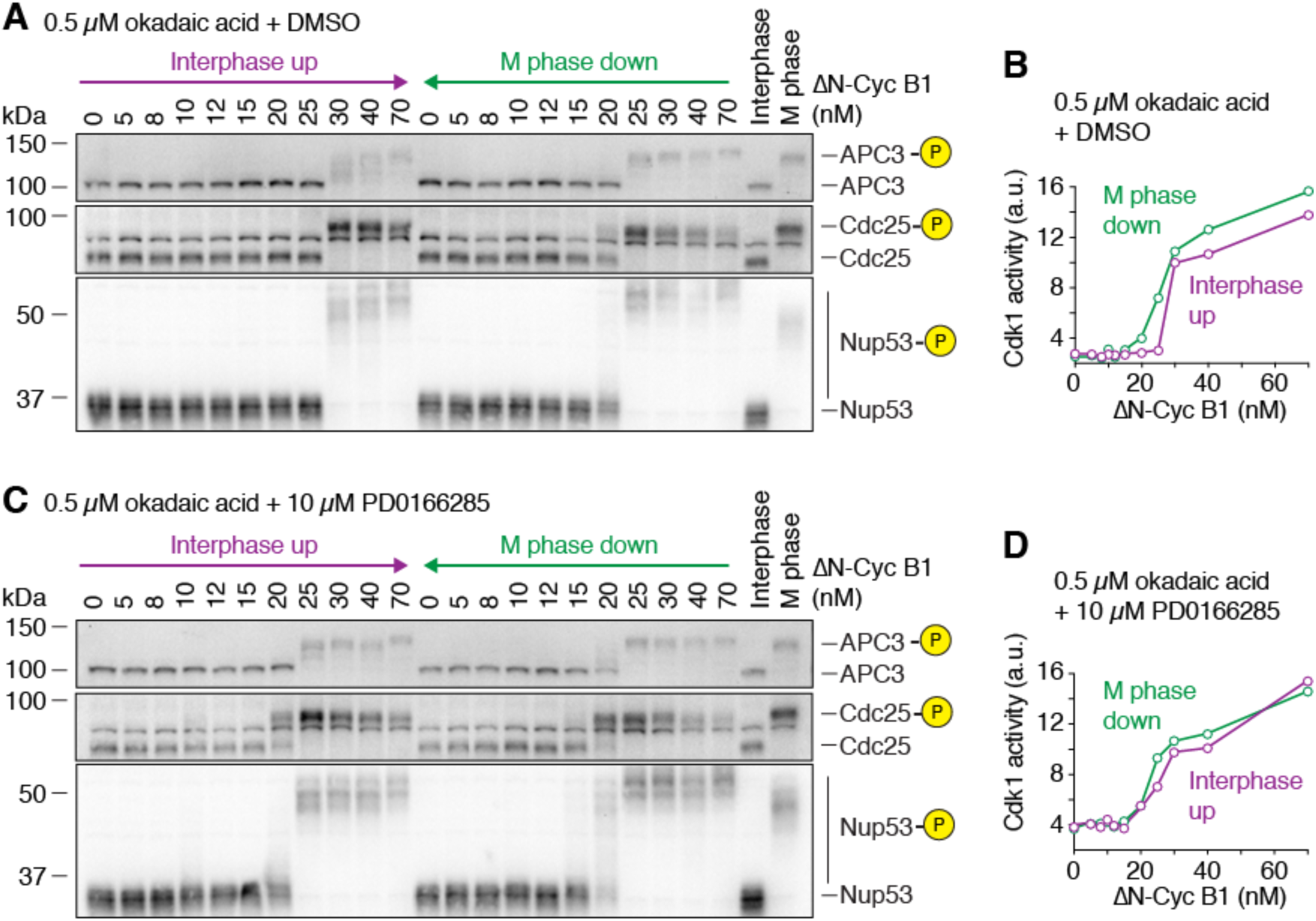
The Cdk1 and PP2A switches both contribute to the bistability. (A-D) The dose-response relationship of the phosphorylation state of several Cdk1 substrates shown in (A) and (C) as well as the Cdk1 activity shown in (B) and (D) was measured in the presence of 0.5 *µ*M okadaic acid and in the absence (A and B) or presence (C and D) of the Wee1/Myt1 inhibitor (10 *µ*M PD0166285). Hysteresis was still detectable if only PP2A activity was partially inhibited, but was almost abolished if Wee1 was inhibited in addition. The measurements were performed in parallel using the same extract. Cdk1 activity is shown as the mean of a technical duplicate with connecting lines.

### Substrate dephosphorylation lags behind Cdk1 inactivation during mitotic exit

In order to investigate how the steady-state properties of Cdk1 and PP2A contribute to the temporal dynamics of substrate phosphorylation, we turned to cycling *Xenopus* egg extracts. These extracts carry out cell cycle transitions homogenously and with good temporal resolution, and thus allow for a detailed dynamic description of when the phosphorylation and dephosphorylation of various Cdk1 substrates occurs.

We first focused on Cdk1 activity and the phosphorylation dynamics of a number of substrates. In the experiment shown in Figures 7A and 7B, at the first time point (45 min into the first mitotic cycle) the cyclin B2 levels were already ∼75% maximal and the Cdk1 activity was ∼50% maximal, but there was no detectable phosphorylation of the 7 mitotic substrates shown. The onset of mitosis occurred at 47-49 min, and was marked by the phosphorylation of all 7 substrate proteins. Phosphorylation was maximal by ∼52 min, with the late substrate APC3 lagging slightly behind the other substrates (Figure 7A). Over the same time interval, cyclin B2 levels were maximal and Cdk1 activity increased 2-fold (Figure 7B). This increase was likely caused by flipping the Cdk1 switch, as Cdc25 hyperphosphorylation and Wee1 T150 phosphorylation became detectable at that time (Figures 7A and 7B). Greatwall and ENSA also became phosphorylated around this time, suggesting that during mitotic entry the Cdk1 and the PP2A-B55 switches are engaged roughly contemporaneously.

**Figure 7.**
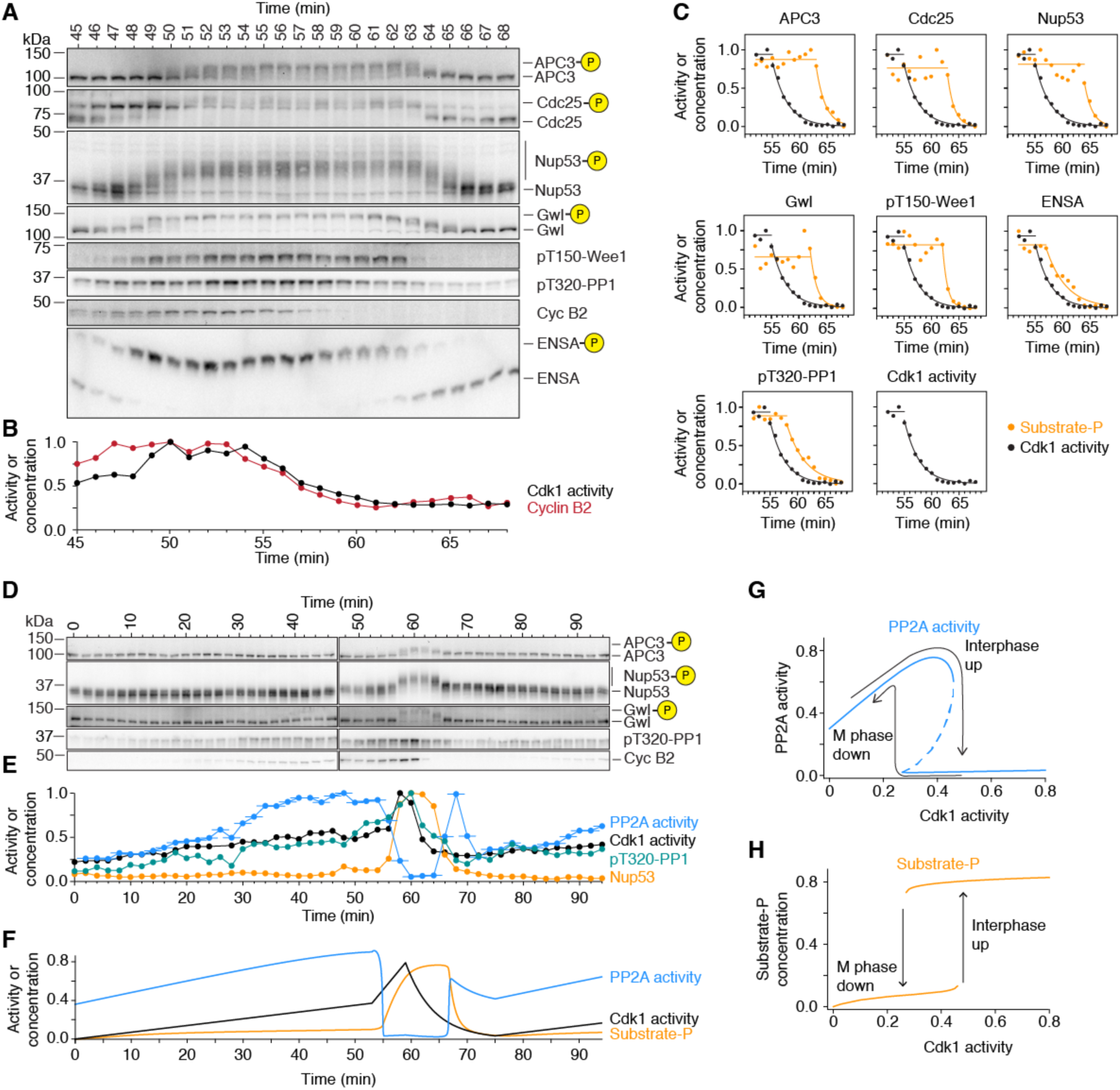
PP2A-B55 activity peaks prior to mitotic entry and during mitotic exit. (A-C) Phosphorylation and dephosphorylation of Cdk1 substrates show a distinct time lag compared to the increase and decrease in cyclin B concentration and Cdk1 activity. The changes in Cdk1 activity ((B) and (C), mean of a technical duplicate), cyclin B2 concentration ((A) and (C)) and the phosphorylation of several substrates (B) were measured with high temporal resolution in a cycling extract progressing through mitosis. Note that cyclin B2 and Cdk1 activity are minimal at 61 min while most substrates are still hyperphosphorylated ((B) and (C), for Cdk1 activity and the phospho-species (as a fraction of total signal) exponential decays were fitted to the declining part of time course). (D and E) PP2A-B55 activity peaks prior to mitotic entry and during mitotic exit. The phosphorylation of two mitotic substrates (D, quantification for Nup53 and PP1 threonine 320 shown in (E) in orange and green, respectively) as well as the concentration of cyclin B2 (D) and the activity of Cdk1 and PP2A-B55 (in (E) black and blue, respectively) were measured in a cycling extract progressing through mitosis. For PP2A-B55 the measurements, the assays were carried out on undiluted extracts; thus, the PP2A-B55 activity could be changing during the 3 min the phosphatase assay is performed. Accordingly, we have plotted the time of each PP2A-B55 measurement as the middle of this incubation period, and show the range of the assay time as a horizontal line. Note that in contrast to PP2A-B55 activity, PP1 threonine 320 phosphorylation closely follows the activity of Cdk1 (D and E). An additional independent experiment is shown in Figures S5A and S5B. (F and G) A simple ODE model incorporating a positive regulation of Cdk1 on PP2A-B55 in addition to the described negative regulation (for details see Figure S6 and the Supplementary text) can describe the temporal dynamic (F) as well as the steady-state response of PP2A-B55 activity (G, blue) and substrate phosphorylation (H, orange) and account for the bistability as well as the biphasic response.

Cyclin B2 levels and Cdk1 activity began to fall at ∼54 min, were half-maximal by ∼57 min, and were back to basal levels by ∼61-62 min (Figures 7B and 7C). In contrast, most of the mitotic substrates did not begin to be dephosphorylated until about 63 min, and then decayed with half-times of ∼2 min (Figures 7A and 7C). PP1 and ENSA appeared to begin being dephosphorylated slightly before the other mitotic substrates at around 57-58 min (Figure 7C). Thus there was a lag of about 6 min between the time when Cdk1 activity had fallen to 50% maximal—an activity that was apparently insufficient to induce substantial substrate phosphorylation during M-phase entry—and when the dephosphorylation of most mitotic substrates began. This suggests that Cdk1 inactivation alone is insufficient to induce detectable substrate dephosphorylation.

### The dynamics of PP2A-B55 activity is a major determinant of the timing of substrate phosphorylation during mitotic entry and exit

We also directly measured PP2A-B55 activity together with cyclin B2 levels, Cdk1 activity, and substrate phosphorylation during mitotic entry and exit. During interphase, as the cyclin B2 concentration and Cdk1 activity gradually rose about 2-fold, PP2A-B55 activity rose by ∼4 fold (Figures 7E and S5B). This is consistent with the finding that sub-threshold levels of Cdk1 activity increase the steady-state activity of PP2A-B55 (Figure 4C). To our knowledge this activation of PP2A-B55 during interphase has not been remarked upon previously.

However, it can be seen in time course data presented by Mochida and Hunt [66]. The increase in PP2A-B55 activity appeared to completely suppress the phosphorylation of APC3 and Nup53 (Figures 7D-E and S5A-B).

At 56 min, PP2A-B55 activity began to fall, and by 60 min it had fallen 17-fold. The decrease in PP2A-B55 was closely followed by an additional ∼2-fold increase in Cdk1 activity and the rapid accumulation of mitotic substrate phosphorylation.

PP2A-B55 activity remained low for the duration of mitosis, but then showed a pronounced spike in activity during mitotic exit. The magnitude of the spike in PP2A-B55 activity was comparable to the PP2A-B55 activity right before mitotic entry, about 14-fold, and coincided with the onset of substrate dephosphorylation. PP2A-B55 activity eventually returned back to its basal interphase level before starting to increase again during the next cycle (Figures 7E and S5B). Thus, PP2A-B55 activity peaks at two different points in the cell cycle, just prior to mitotic entry and during mitotic exit. Moreover, the dynamics of PP2A-B55 activity appear to play a large role in determining substrate phosphorylation; during entry, a small increase in Cdk1 activity (∼2-fold) is able to cause a large change in substrate phosphorylation because PP2A-B55 activity decreases ∼17 fold, and during exit, the large change in Cdk1 activity due to cyclin proteolysis is unable to cause substrate dephosphorylation until PP2A-B55 activity increases sharply.

### Bistability plus incoherent feedforward regulation can account for the relationship between Cdk1 activation, PP2A-B55 activity, and substrate phosphorylation

Given the complexity of the PP2A-B55 regulatory system, it is difficult to simply intuit whether the system as currently understood is able to give rise to the observed bistability and biphasic responses in the steady-state regulation of PP2A-B55, and the observed dynamics of substrate phosphorylation during mitotic entry and exit. To clarify this point, we constructed a relatively simple model of the PP2A-B55 system that included two key circuit motifs: incoherent feedforward regulation of PP2A-B55 by Cdk1, and double-negative feedback between PP2A-B55 and the ENSA and Arpp19 proteins (considered indistinguishable for the purpose of the model and hereafter referred to as ENSA).

The core of the model is a set of 4 ordinary differential equations (ODEs) and two conservation equations (yielding a net of 2 independent ODEs), shown schematically in Figure S6A and in detail in the supplementary text, that describe the interplay between ENSA, phosphorylated ENSA, and PP2A-B55. For this model, we assumed that Gwl activity exhibits an ultrasensititive but not bistable response to Cdk1 activity. In such a model bistability arises from the interaction of phosphorylated Arpp19/ENSA and PP2A-B55. Theoretical considerations show that a stoichiometric inhibitor which is also inactivated by the enzyme it inhibits cannot, in the simplest models, produce bistability [70, 71]. However, small modifications of the simplest model can allow for bistability [71]. Therefore, we implemented one of these: We assumed that phosphorylated Arpp19/ENSA interacts with PP2A-B55 in two ways –stochiometrically inhibiting PP2A-B55 via one binding site and being dephosphorylated by PP2A-B55 via a different binding site (see details in the supplementary text). Note that this is an ad hoc assumption; so far there is no direct evidence for two such binding modes. Nevertheless, this model was able to generate a bistable response for the relationship between the activity of Cdk1 and the concentration of free PP2A-B55 (Figures S6B and S6C).

The second part of the model aimed to account for the biphasic nature of PP2A-B55 regulation. We assumed that Cdk1 increased the activity of free PP2A-B55 above its basal level. The mechanism is unknown; for simplicity we modeled this positive regulation as direct phosphorylation of PP2A-B55 by Cdk1 although we speculate that in reality this interaction is more indirect. In addition, we assumed that Cdk1 exerted a negative effect on PP2A-B55 by stimulating, through Gwl, the phosphorylation of ENSA. This type of signaling motif, where an upstream protein (Cdk1) exerts opposite effects on a downstream protein (PP2A-B55) through two different pathways, is referred to as incoherent feedforward regulation [72], and it is one way of generating biphasic responses.

We manually scanned the model’s parameters and were able to identify parameters that reproduced both the bistable, biphasic steady state response of PP2A-B55 activity and substrate phosphoryaltion (Figures 7) and the dynamics of PP2A-B55 regulation and substrate phosphorylation (Figures 7A–7F). As observed experimentally, substrate phosphorylation remained low during the time when Cdk1 and PP2A-B55 activity increased concurrently, rapidly increased when Cdk1 activity increased 2-fold and PP2A-B55 activity decreased about 20-fold, and stayed high until PP2A-B55 activity peaked during mitotic exit (Figure 7F). To recreate the peaks in PP2A-B55 activity during mitotic entry and exit seen in the time course data (Figures 7E and S5B, blue curves), it was necessary to assume that the regulation of PP2A-B55 by Cdk1 was fast compared to the changes in Cdk1 activity. In fact, this appears to be the case, given that the relationship between PP2A-B55 activity and Cdk1 activity seen in time course experiments is similar to that expected from the steady-state response curves (Figures S5C and S5D). However, the modeled dynamics of the PP2A-B55 peak during mitotic exit still do not exactly match the experimental data. The reasons for this discrepancy remain to be determined.

Thus, a simple incoherent feedforward model with double-negative feedback and stoichiometric inhibition can qualitatively account for the bistable, biphasic steady-state activity of PP2A-B55 and the dynamics of PP2A-B55 activation and substrate phosphorylation during mitotic entrance and exit. Presumably the details of the phosphatase dynamics that are not accounted for by the two ODE model would require a better understanding of the biochemistry of this system and a more complicated model.

## Discussion

Here we have investigated the steady-state and temporal behavior of mitotic substrate phosphorylation, and the roles of Cdk1, PP1, and PP2A-B55 in generating the dramatic events of mitotic entry and exit. We found that even when Cdk1 activity is forced to be a graded, monostable function of the cyclin B1 concentration, several mitotic phosphoproteins still exhibit bistable responses (Figure 1), including the critical mitotic regulator APC/C (Figure 2). Thus, even though the Wee1/Cdc25/Cdk1 system is essentially inoperative during embryonic cycles 2-12 [37–39], the Cdk1 system as a whole still includes a bistable trigger and can still be viewed as a relaxation oscillator—an oscillator built from toggle switches. This class of oscillator is common in biology, and, compared with oscillator circuits that are not built out of switches, tends to be especially robust and tunable [73]. The bistability can be attributed to PP2A-B55: its regulators ENSA and Arpp19 exhibit bistable phosphorylation, and PP2A-B55 (but not Gwl) exhibits bistable activity changes (Figures 4, S2, and S3). In addition, perturbing PP2A activity with low concentrations of okadaic acid abolished this bistability (Figures 5 and 6). These findings agree well with *in vitro* reconstitution studies of the *Xenopus* PP2A-B55/Gwl system [33] and provide a molecular mechanism to explain why it takes lower concentrations of a Cdk1 inhibitor to block mitotic entry than to induce mitotic exit [34]. In contrast, there was no evidence for bistability in the response of PP1 (Figure 3). Instead, PP1 T320 phosphorylation closely tracked Cdk1 activity (Figure 7), suggesting that PP1 activity simply varies inversely with Cdk1 during mitotic entry and exit.

In addition, we found that Cdk1 activity at sub-threshold levels activates PP2A-B55. This is apparent in both the steady-state response of PP2A-B55 (Figures 4 and S2) and in the time course studies (Figure 7 and S5). This incoherent feedforward regulation has a number of significant consequences for the systems-level behavior:

1. Because PP2A-B55 activity gradually increases in parallel to Cdk1 activity when approaching mitosis, the kinase-to-phosphatase ratio remains low and substrates remain dephosphorylated during this time. Indeed, no mitotic substrate phosphorylation is detectable until PP2A-B55 is inactivated (Figures 7 and S5). It has been pointed out that the reciprocal regulation of Cdk1 and its counteracting phosphatases avoids energy-intensive futile cycles of phosphorylation and dephosphorylation [60]. While this energy-saving measure may be beneficial when keeping the system in a stable state, by itself it would render the system slower to respond [74]. Up-regulating the phosphatase in addition to the kinase activity prior to mitotic entry increases futile cycling and the flux in the system, without changing the net phosphorylation, which allows for a rapid and abrupt transition from hypo-to hyperphosphorylation once the phosphatase is inactivated. Consistent with this idea, mitotic entry has recently been associated with high energetic costs during the embryonic cell cycle of zebrafish [75].
2. Whereas the difference between the basal and mitotic activities of PP2A-B55 are 4- to 5-fold, the additional activation of PP2A-B55 during the ramp up to mitosis results in an overall 15- to 20-fold change in PP2A-B55 activity at the time of mitotic entry. Note that by comparison, the activity of Cdk1 changes in a relatively subtle fashion during mitotic entry, with the activity increasing about 2 fold in *Xenopus laevis* (here and Ref. [11, 76–78] and other organisms as well [79, 80]). Thus, the changes in PP2A-B55 appear to be the main determinants of the quantitative changes in substrate phosphorylation during mitotic entry.
3. The spike in PP2A-B55 activity during mitotic exit ensures that substrates are rapidly dephosphorylated. Indeed, although there is a lag between the time when Cdk1 falls to sub-mitotic levels and substrate dephosphorylation commences, once it begins, substrate dephosphorylation is essentially complete within 2-3 min (Figure 7). This allows substrate phosphorylation to be reversed rapidly and abruptly during mitotic exit. The mechanism of activation is unknown, but we suspect that it is indirect. This is based on the fact that such an activation has not been observed in in vitro reconstitution assays [33]. Direct phosphorylation of threonine 304 of the catalytic subunit of the PP2A-B55 trimer by Cdk1 has recently been reported in human cells in cell culture [31]. However, this phosphorylation has been implicated in a negative rather than positive regulation of PP2A-B55 activity by destabilizing the trimeric complex during mitosis [31, 81]. In *X. laevis* egg extracts, on the other hand, cell cycle dependent changes of PP2A-B55 complex formation have not been observed [6].

PP2A-B55 is thought to directly dephosphorylate Wee1 and Cdc25 [6], thereby negatively regulating Cdk1 activity via tyrosine-15 phosphorylation. This raises the question how such a system could switch into a state of high Cdk1 activity if Cdk1 steadily upregulates PP2A-B55. One explanation could be that an increase in the basal activity of Y15-phosphorylated cyclinB-Cdk1 complex due to increasing cyclin B levels suffices for flipping the switch. Such an increase is consistent with the slow but steady increase in Cdk1 activity during interphase that we observe in Figure 7E and with previous reports from human cells (e.g., [9]). Other activities might also contribute to the flipping of the switch, for example cyclin A-Cdk1 which is not inhibited by Wee1/Myt1 [82].

Note that during both mitotic entry and mitotic exit, the changes in Cdk1 appear to be permissive, whereas the changes in PP2A-B55 activity appear to be instructive. Nevertheless, in the *Xenopus* system, Cdk1 does appear to be the master regulator of M-phase; cyclin accumulation drives Cdk1 activation, and then Cdk1 activation drives PP1 and PP2A-B55 inactivation. However, in other systems, it is possible that the trigger for mitosis could initially act upon any of these components—the central players Cdk1, PP1, and PP2A-B55, or the intermediaries Wee1, Cdc25, Gwl, and ENSA/Arpp19—since they are all interconnected by feedback loops.

## Acknowledgments

We would like to thank Wolfram Antonin, Satoru Mochida, Andreas Heim, Thomas Mayer, Qiong Yang, Matthew Swaffer and Connie Phong for reagents and helpful discussions, Renping Qiao Coudevylle and Jan Michael Peters for supplying purified non-degradable cyclin B1, Satoru Mochida and Bela Novak for sharing their manuscript prior to publication and Silke Hauf, Andreas Boland, Oshri Afanzar, Xianrui Cheng, William Huang, Connie Phong, and Ivan Zheludev for thoughtful comments on the manuscript. The work was supported by grants from the National Institutes of Health (R01 GM046383 and P50 GM107615, J.E.F.) and a postdoctoral fellowship from the German Research Foundation (KA 4476/1-1, J.K.).

## Author Contributions

J.K. and J.E.F. conceived the project; J.K. performed all experiments; J.K., L.G., and J.E.F. analyzed the data and carried out the modeling; J.K. and J.E.F. wrote and all authors edited the manuscript.

## Methods

### Egg extract preparation

Eggs were collected from female *Xenopus laevis* frogs 16-18 hours after egg laying was induced by injection of 500 U of human chorionic gonadotropin (Chorulon from Merck, Sigma #CG10-10VL). Cytostatic factor-(CSF-) arrested and cycling extracts were prepared as described previously [83], except that for the cycling extracts eggs were activated with the calcium ionophore A23187 (200 ng/µL, Sigma Aldrich # C7522) rather than electric shock. CSF extracts were stored at -80°C until use; cycling extracts were used directly after preparation.

Animal work was conducted in accordance with relevant national and international guidelines and all animal protocols were approved by the Stanford University Administrative Panel on Laboratory Animal Care.

### Titration of non-degradable cyclin B1

CSF extracts were thawed at 20°C for 5 min. Extracts were supplemented with cycloheximide (100 *µ*g/mL) and where applicable with the Wee1/Myt1 inhibitor PD0166285 (Abcam, ab219507) or okadaic acid (EMD Millipore, #495609). The extract was then treated in one of two ways: (1) Aliquots of the extract were supplemented with different concentrations of recombinant non-degradable cyclin B1 (ΔN-Cyc B1) purified from insect cells, incubated for 1 h at 20°C and then supplemented with CaCl_2_ to a final concentration of 0.8 mM and incubated for another hour (‘M phase down’) or (2) aliquots of the extracts were first treated with 0.8 mM CaCl_2_ for 1 h at 20°C and then supplemented with different concentrations of non-degradable cyclin B1 and incubated for another hour (‘Interphase up’). All samples were mixed regularly during the incubation.

### Immunoblotting and antibodies

Protein samples were resolved on 10% Criterion Tris-HCl precast gels (Biorad, #3450011) or in the case of the phosphorylated species of Arpp19 and ENSA on a 10% Tris-HCl SDS-PAGE supplemented with 10 μM Phos-tag reagent (NARD Institute, AAL-107) and then transferred onto PVDF membranes. The following antibodies were used for detection of the respective proteins: mouse *α*-Cdc27 (BD Biosciences, #610455), mouse *α*-cyclin B2 (Santa Cruz Biotechnology, #sc-53239), rabbit *α*-Nup53 serum [84], rabbit *α*-PPP1A pT320 (Abcam, #ab62334), rabbit *α*-Cdc25C [44], rabbit *α*-Wee1 pT150 [85], rabbit *α*-Greatwall serum [60], rabbit *α*-ENSA serum [60], rabbit *α*-Arpp19 [63], and rabbit *α*-Cdk1 pY15 (Cell Signaling Technology, #9111L). For the detection of the primary antibody, peroxidase-linked sheep *α*-mouse IgG (GE Healthcare, #NA931V) and AMDEX goat *α*-rabbit IgG (GE Healthcare, RPN4301) was used. Chemiluminescence was detected on a BioRad ChemiDoc MP Imaging system using SuperSignal West Maximum Sensitivity Substrate (ThermoFisher Scientific, #34095) or Immobilon Western Chemiluminescent HRP Substrate (EMD Millipore, #WBLKS0500).

### H1 kinase activity assay

H1 kinase assays followed the previously described protocol [83]. In brief, 2 *µ*L frozen extract sample were diluted in 98 *µ*L EB buffer (80 mM *β*-glycerophosphate, 20 mM EGTA, 15 mM MgCl_2_, pH 7.4). Ten microliters of diluted extract were mixed with 10 *µ*L reaction buffer (20 mM HEPES pH 7.5, 5 mM EGTA, 10 mM MgCl_2_, 200 mM ATP, 10 *µ*g histone H1 (Millipore, #14-155), 20 *µ*M PKA inhibitor IV (Santa Cruz Biotechnology, #sc-3010) and 2.5 *µ*Ci [*γ*-^32^P]-ATP) and immediately incubated for 3 min at 20°C. The reaction was stopped by adding 20 *µ*L 3x SDS PAGE loading dye. Five microliters of each sample were run on a 10% Criterion Tris-HCl precast gel (Biorad, #3450011), transferred onto a PVDF membrane and dried. The radiolabelled histone H1 was then detected using a BAS Storage Phosphor screen (GE Healthcare) and read out using a Phosphorimager Typhoon 8600 (Molecular Devices).

### APC/C activity assay

APC/C activity assays were performed as described previously [45]. Briefly, securin-CFP was *in vitro* translated using the TNT SP6 high-yield wheat germ protein expression system (Promega, #L3261). Three or 4 *µ*L of in vitro translated securin-CFP was added to 50 or 80 *µ*L extract. The extract was split into aliquots of 20 *µ*L and added to a black 384-well plate (Greiner, #781076). Securin-CFP signal was followed using a plate reader (Molecular Devices Flexstation II). The degradation rate was calculated by normalizing the data to the starting value and the background control and fitting a single exponential decay. All measurements were performed in duplicate or triplicate.

### PP2A-B55 activity assay

PP2A-B55 activity assays were performed as previously described [66] with slight modifications. In brief, maltose-binding protein fused to amino acids 38 to 62 of *Xenopus laevis* Cdc20L (Fizzy) was recombinantly expressed in *E. coli* and affinity purified using amylose resin. Without elution, about 1 mg of protein was phosphorylated in 200 *µ*L kinase reaction buffer (20 mM HEPES pH 7.7, 10 mM MgCl_2_, 15 mM KCl, 1 mM EGTA, 5 mM NaF, 20 mM *β*-glycerophosphate, 10 *µ*M ATP, 2.5 *µ*g cyclin A2/CDK2 (Sigma, #C0495), 60 *µ*Ci [*γ*-^32^P]-ATP) overnight at 37°C. The resin was extensively washed and the labeled substrate eluted with elution buffer (20 mM Tris-HCl pH 7.5, 150 mM NaCl, 10 mM maltose). The substrate was concentrated and stored at -20°C until use.

For measuring PP2A-B55 activity, 1 *µ*L of substrate (>15,000 cpm) was added to 5 *µ*L of extract and incubated for 12 min at 20°C (3 min for the time course measurements). The reaction was stopped by adding 20 *µ*L 10% ice-cold TCA and stored on ice until further processing. Samples were spun for 10 min at 14,000 g and 20 *µ*L of the supernatant was transferred to a fresh tube. Thirty microliters of 5% ammonium molybdate in 0.5 M sulfuric acid was added and mixed. Fifty microliters of water-saturated heptane/butanol was added and the solution vortexed for 30 sec. The solution was spun for 10 min at 14,000 g and 30 *µ*L of the organic upper phase was used for detection of the inorganic phosphate using a scintillation counter.

### Greatwall kinase activity assay

Greatwall kinase activity assays were performed following the previously described protocol [63]. In brief, 1 *µ*L egg extract was diluted in 16.5 *µ*L reaction buffer (80 mM *β*-glycerophosphate, 20 mM EGTA, 15 mM MgCl_2_, 10 uM okadaic acid, 1x protease inhibitor (Roche, cOmplete #11873580001), 100 *µ*M ATP, 10 *µ*M PKA inhibitor IV (Santa Cruz Biotechnology, #sc-3010), 120 nCi/uL [*γ*-^32^P]ATP, 0.05 *µ*g/µL Arpp19). The reaction was incubated at 20°C for 3 min and stopped by adding 17.5 *µ*L of 3x SDS loading dye. Five microliters of each sample were run on a 4-20% Criterion Tris-HCl precast gel (Biorad, #3450034), transferred onto a PVDF membrane and dried. The radio-labelled Arpp19 was then detected using a BAS Storage Phosphor screen (GE Healthcare) and read out using a Phosphorimager Typhoon 8600 (Molecular Devices). Alternatively, Gwl was first immunoprecipitated from 5 *µ*L of extract using rabbit *α*-Greatwall serum [60] coupled to magnetic Protein G beads (Invitrogen, #10004D) for 20 min at 20°C under shaking. The beads were washed three times with PBS and all supernatant was discarded. Gwl kinase activity was then measured as described above except that the beads were resuspended in 16.5 *µ*L reaction buffer that did not contain okadaic acid.

### Purification and thiophosphorylation of Arpp19

Purification and thiophosphorylation of Arpp19 were performed as described previously [63] with slight modifications. In brief, N-terminally His-tagged Arpp19 (*X. laevis*) was overexpressed and purified from *E. coli* BL21(RIL) via affinity purification. mRNA encoding 3xFlag-Xl-Gwl (K71M, full length) was transcribed in vitro using an mMESSAGE mMACHINE kit (Invitrogen, #AM1344), poly(A)tailed using a Poly(A) tailing kit (Invitrogen, #AM1350), and purified using the MEGAclear Transcription Clean-Up kit (Invitrogen, #AM1908).

Freshly prepared CSF extract (200 *µ*L) was supplemented with 2.5 *µ*M okadaic acid, 1.4 *µ*M non-degradable cyclin B (ΔN90-cyclin B from sea urchin) and 8 *µ*g Gwl K71M mRNA in order to translate and phosphorylate 3xFlag-Gwl K71M. The translated protein was immunoprecipitated using 1.2 mg of anti-Flag coupled magnetic beads (Sigma-Aldrich, #F1804, covalently bound to Protein G beads (Invitrogen, #10004D)). Beads were washed 3 times briefly and 3 times for 5 min with CSF-XB buffer (10 mM HEPES pH 7.7, 100 mM KCl, 1 mM MgCl_2_, 0.1 mM CaCl_2_, 50 mM sucrose, 0.5 mM EGTA, 1 mM MgCl_2_). Full-length Arpp19 (100 *µ*g) in phosphorylation buffer (10 mM Tris-HCl pH 7.4, 10 mM KCl, 30 mM NaCl, 0.5 mM EGTA, 20 mM beta-glycerophosphate, 5 mM MgCl_2_, 1 mM DTT, 1 mM MnCl_2_, 30 nM okadaic acid, 1 mM ATP-γ-S) was added to the beads and incubated overnight at 30°C. Subsequently, the beads were removed and the supernatant dialyzed against storage buffer (20 mM HEPES pH 7.7, 150 mM KCl, 10% glycerol, 1 mM DTT).

## Supplemental Information

### Supplemental figures

**Figure S1, related to Figure 1.**
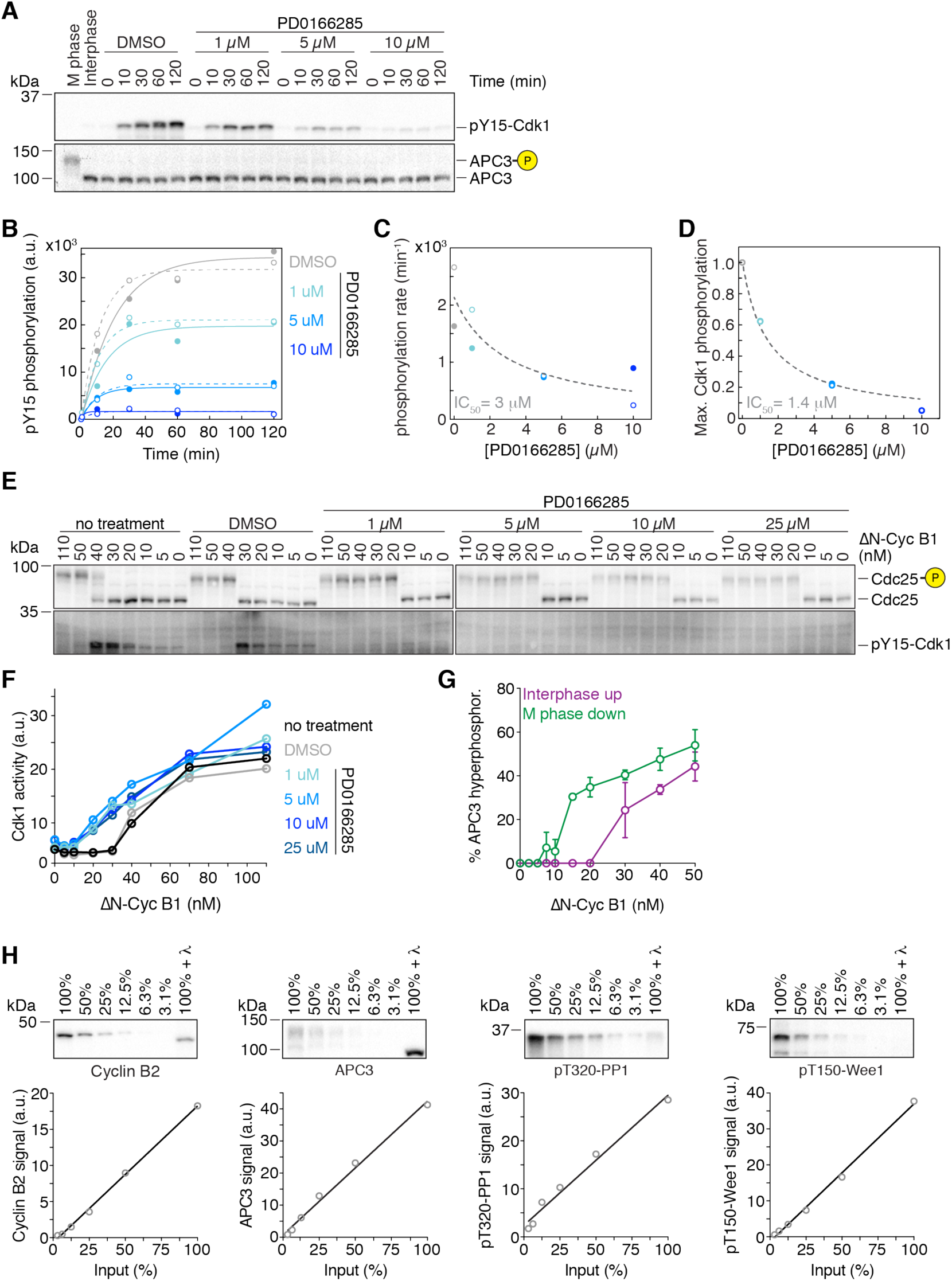
Characterization of the Wee1/Myt1 inhibitor PD0166285 and the antibodies used in this study. (A) Titration of the Wee1 inhibitor PD0166285. An interphase extract supplemented with cycloheximide was treated with different concentrations of PD016885 or DMSO. Because Wee1 only phosphorylates the Cdk1-cyclin B complex, complex formation was initiated by adding non-degradable cyclin B. To prevent the active Cdk1-cyclin B complex from inhibiting the Wee1 kinase, the Cdk1 inhibitor roscovitine (1 mM) was added to the extract. The phosphorylation of Cdk1 was followed over time using a phospho-specific antibody recognizing Cdk1 tyrosine 15 phosphorylation. APC3 functions as a loading control and as a control for low Cdk1 activity. (B) Quantification and exponential fits, full circles and lines respectively, of the experiment shown in (A), and an additional technical replicate (clear circles and dashed lines). (C) The apparent first-order phosphorylation rate of Cdk1 tyrosine 15 by Wee1 kinase (derived from the fits in (B)) as a function of the concentration of the Wee1/Myt1 inhibitor PD0166285. The fit (dashed line) yielded an IC_50_ of 3 *µ*M. (D) Maximal phosphorylation signal (normalized to the DMSO control) after 120 min of incubation plotted as a function PD0166285 concentration. The fit (dashed line) yielded an IC_50_ of 1.4 *µ*M. (E and F) Dose-response of Cdc25 hyperphosphorylation and Cdk1 tyrosine 15 phosphorylation (E) as well as Cdk1 activity (F) as a function of non-degradable cyclin B1 (ΔN-Cyc B1) for different concentrations of the Wee1 inhibitor PD0166285 during M phase entry. Inhibitor concentrations of 5 *µ*M and higher resulted in a lower threshold for Cdc25 hyperphosphorylation, absence of any detectable Cdk1 tyrosine 15 phosphorylation (E), and a more gradual increase in Cdk1 activity (F). (G) Quantitation of the APC3 hyperphosphorylation as a function of ΔN-Cyc B1 concentration. Shown is the mean and the standard error of the mean from 3 independent experiments. (H) Linearity tests for the antibodies used quantitatively in this study. Proteins were detected with the respective antibodies from a dilution series of an M phase extracts as well as from a sample treated with λ phosphatase; signal intensities were quantified and plotted. Solid lines show the linear fit of the data.

**Figure S2, related to Figure 4.**
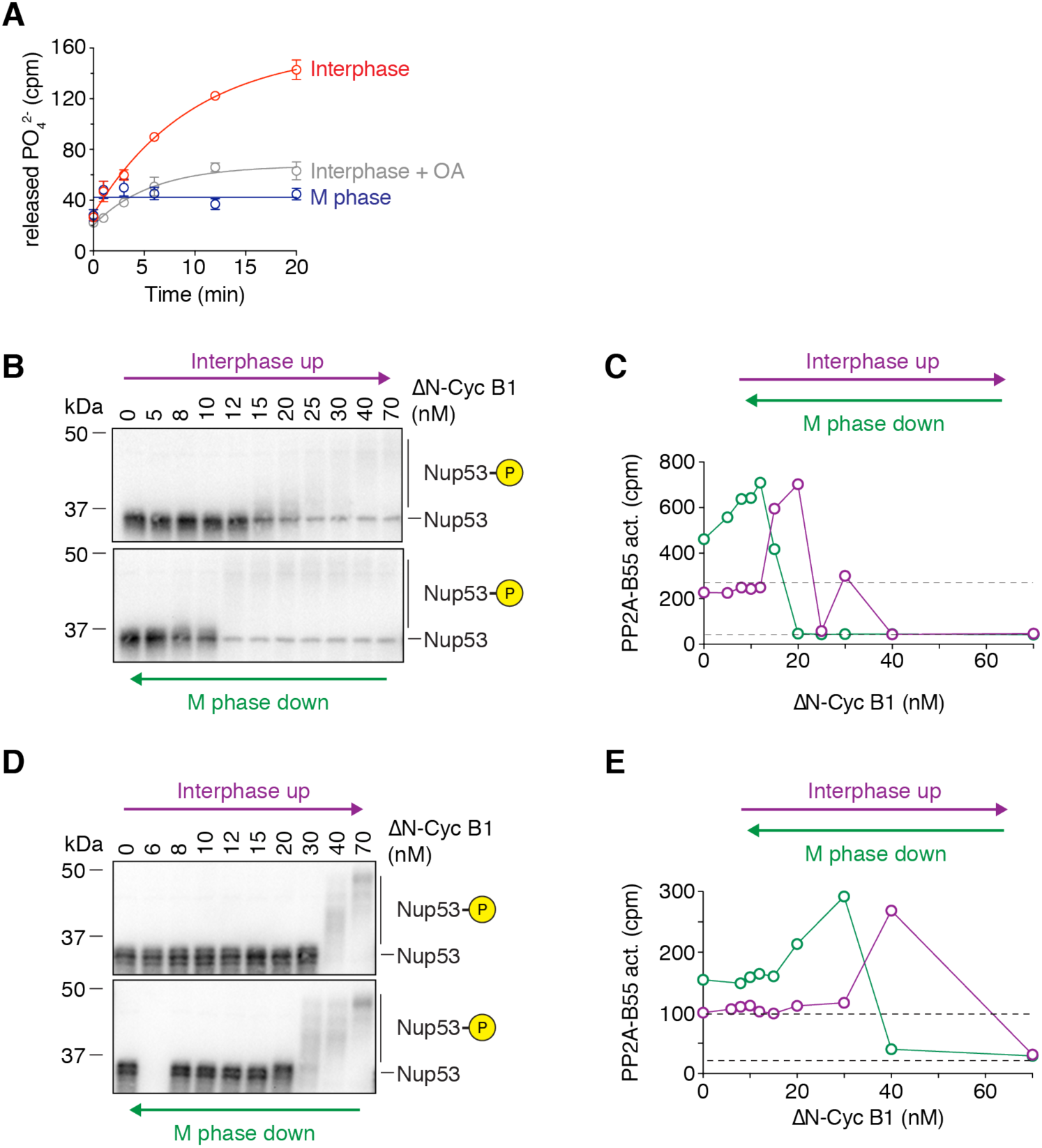
PP2A-B55 activity exhibits bistability. (A) Measuring PP2A-B55 activity. The release of inorganic phosphate from a radioactively labeled PP2A-B55 substrate (MBP-Fzy S50) was followed over time using a scintillation counter. The activity of PP2A-B55 was significantly higher in an interphase extract than in an M phase extract and was inhibited by the addition of 2.5 *µ*M okadaic acid. Shown are the mean and standard deviation of a technical triplicate (circles with error bars) and fitted to either exponentials (interphase, interphase + OA) or a straight line (M phase). (B-E) Two additional independent experiments similar to the one shown in Figures 4B and 4C measuring mitotic substrate phosphorylation (monitored by the mobility shift of Nup53, (B) and (D)) and PP2A-B55 activity ((C) and (E)) as a function of non-degradable cyclin B1 (ΔN-cycB1) approaching steady-state starting from either a state of high (M phase down, green) or low (Interphase up, purple) Cdk1 activity/cyclin B concentration (mean of a technical duplicate, circles, with connecting lines) in the presence of 10 *µ*M PD0166285.

**Figure S3, related to Figure 4.**
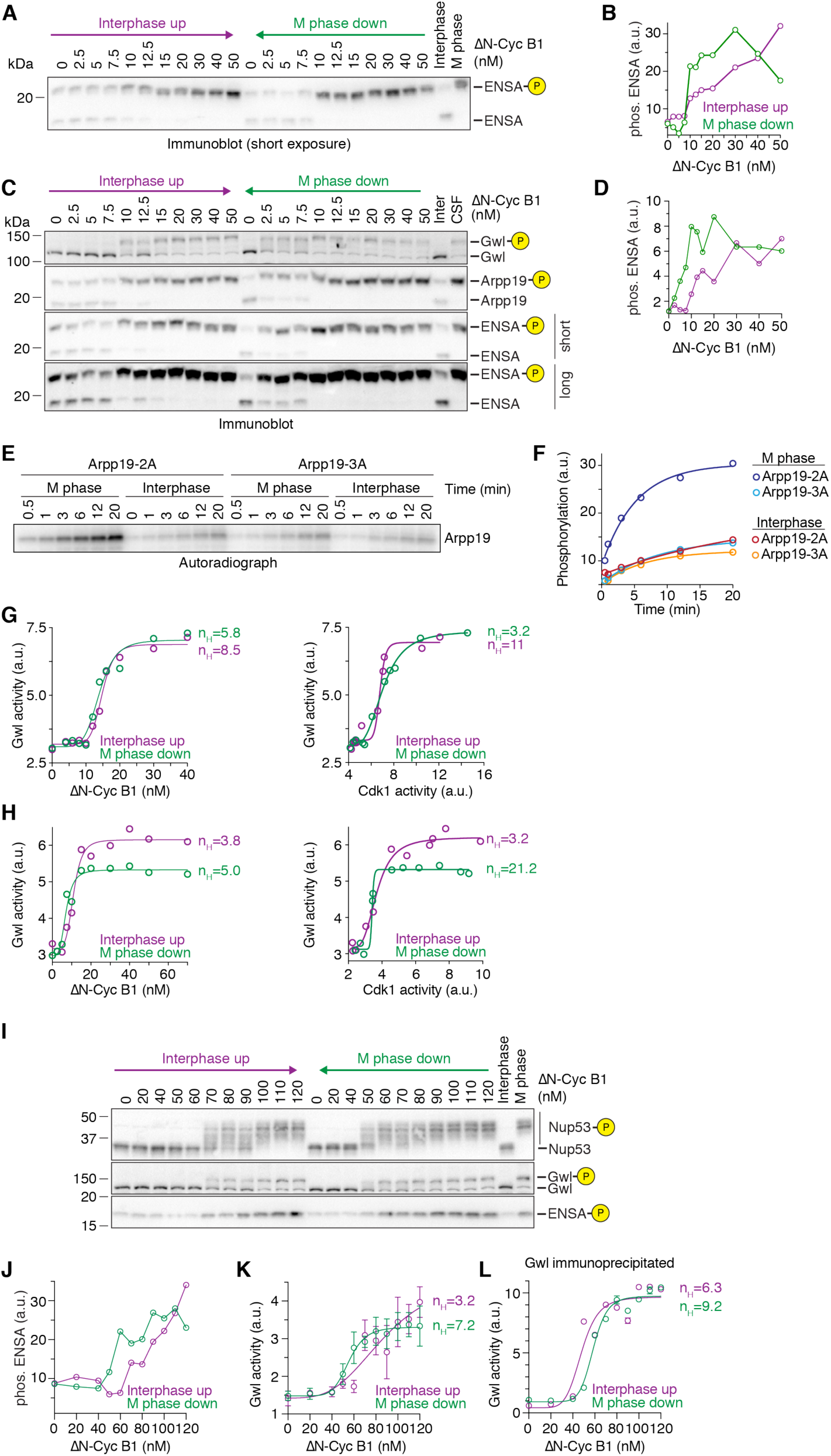
Gwl kinase activity exhibits ultrasensitivity but not bistability. (A) Shorter exposure of the ENSA immunoblot shown in Figure 4D. (B) Quantification of the hyperphosphorylated form of ENSA from the experiment shown in Figure S3A. (C) An additional independent experiment analyzing the dose-response of Greatwall kinase, Arpp19 and ENSA phosphorylation as a function of ΔN-Cyc B1 concentration (similar to Figure 4D). An antibody raised against a conserved region of ENSA (third and fourth blot from the top) and an antibody raised against full-length Arpp19 (second blot from the top) showed similar behavior. Phos-tag (10 *µ*M) was used to enhance the mobility shift upon ENSA and Arpp19 phosphorylation (second to fourth blot from the top). (D) Quantification of the hyperphosphorylated form of ENSA from the experiment shown in Figure S3C. (E and F) Measuring Greatwall (Gwl) kinase activity. Recombinant Arpp19-2A (S28A S109A), which maintains the Gwl phosphorylation site (S67) but carries mutations in two other prominent non-Gwl phosphorylation sites (Cdk1 for S28 and PKA for S109, respectively), or the recombinant Arpp19-3A (S28A S67A S109A), were incubated with M-phase or interphase extract and the incorporation of ^32^P was monitored over time by autoradiography after resolving the protein on an SDS polyacrylamide gel (E). The kinase activity toward Arpp19-2A was increased in M phase. Quantification of the signals from the autoradiograph as a function of time, plus an exponential fit of the data (lines), are shown in (F). (G and H) Two additional independent experiments measuring Gwl activity as a function of non-degradable cyclin B1 (ΔN-Cyc B1, left) or Cdk1 activity (right). The dose-response relationships of Greatwall kinase activity show strong ultrasensitivity but no hysteresis. Fitted Hill curves yielded apparent Hill exponents of 3 to 21. (I-L) An additional independent experiment analyzing the dose-response of Greatwall phosphorylation and activity, and Arpp19/ENSA phosphorylation, as a function of ΔN-Cyc B1 concentration. Arpp19/ENSA phosphorylation was detected using an antibody against the phosphorylated epitope surrounding Ser 62 and Ser 67, respectively. A quantification of the detected phosphorylation signal is shown in (J). Gwl kinase activity was measured directly from the sampled extracts in the presence of okadaic acid (K) similar to (G) and (H), or Gwl was first immunoprecipitated from the extract and Gwl activity was subsequently measured in the absence of okadaic acid (L). Fitted Hill curves yielded apparent Hill exponents of 3 to 9.

**Figure S4, related to Figure 5.**
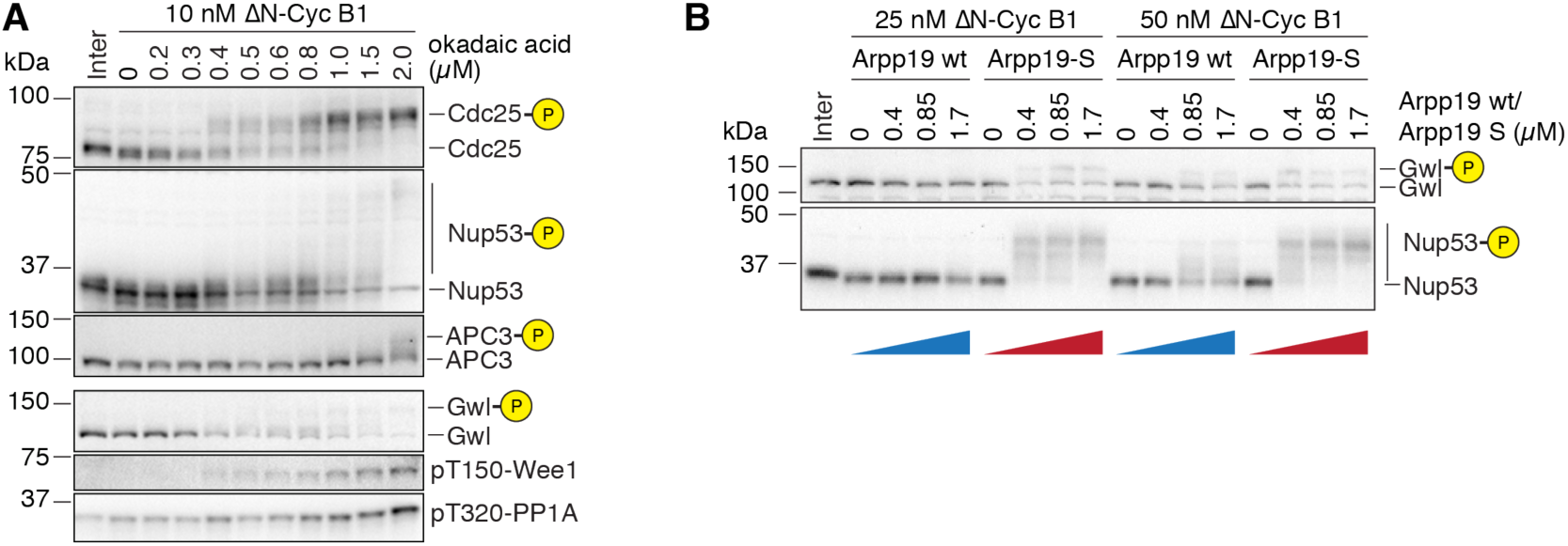
Characterization of the PP2A inhibitors okadaic acid and thio-phosphorylated Arpp19. (A) Titration of the PP2A inhibitor okadaic acid. An interphase extract was supplemented with a low concentration of non-degradable cyclin B (ΔN-Cyc B1) and different concentrations of okadaic acid. After reaching steady-state, the phosphorylation states of several Cdk1 substrates were analyzed by mobility shift. Concentrations between 0.4 *µ*M and 0.6 *µ*M okadaic acid subtly impacted the phosphorylation state of several Cdk1 substrates including Cdc25, Nup53, Gwl and Wee1, but not the phosphorylation of PP1 at threonine 320 and the hyperphosphorylation of APC3. (B) Titration of the PP2A-B55 inhibitor Arpp19-S (thiophosphorylated Arpp19). An interphase extract was supplemented with low concentrations of non-degradable cyclin B (ΔN-Cyc B1) and different concentrations of Arpp19-S. After reaching steady-state, the phosphorylation states of several Cdk1 substrates were analyzed by mobility shift. 0.4 *µ*M Arpp19-S but not unphosphorylated Arpp19 changed the phosphorylation state of both Gwl and Nup53.

**Figure S5, related to Figure 7.**
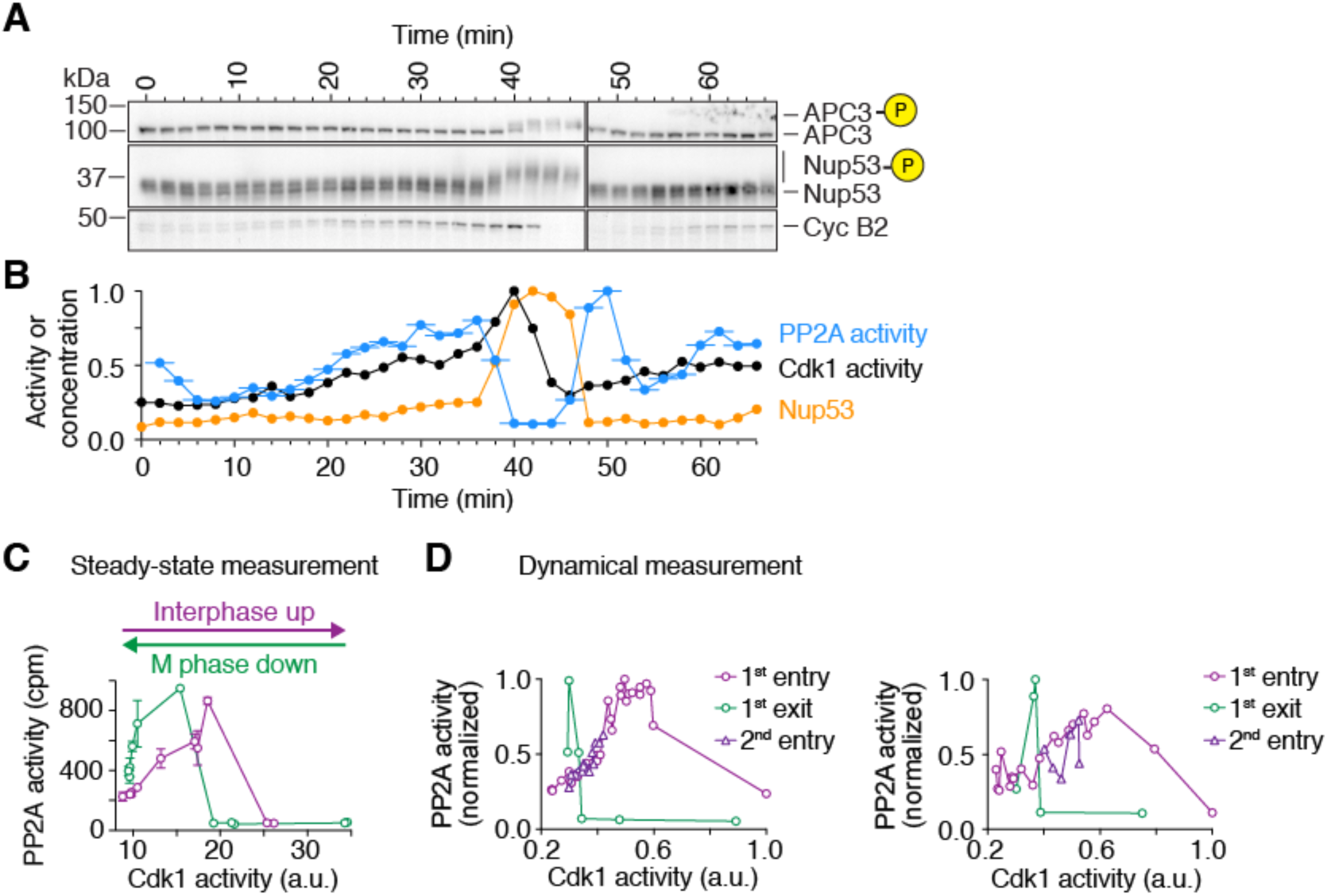
PP2A-B55 activity peaks prior to mitotic entry and during mitotic exit. **(A and B)** An additional time course experiment similar to the one shown in Figure 7D and 7E. Mitotic substrate phosphorylation ((A), quantified for Nup53 in orange in (B)), cyclin B2 concentration (A) and Cdk1 and PP2A-B55 activity ((B), black and blue respectively) were measured every two minutes in a cycling *Xenopus laevis* egg extract. For PP2A-B55 the measurements, the assays were carried out on undiluted extracts; thus the PP2A-B55 activity could be changing during the 3 min if the phosphatase assay. Accordingly, we have plotted the time of each PP2A-B55 measurement as the middle of this incubation period, and show the range of the assay time as a horizontal line. **(C and D)** The steady-state response of PP2A-B55 as a function of Cdk1 activity compared to the relationship between PP2A-B55 activity and Cdk1 activity derived from the time course experiments (Figures 7D, 7E and S5B). The similarity between the steady-state (C) and the time resolved data (D) argues that the regulation of PP2A-B55 activity is fast and close to steady-state at any given time. The steady-state response of PP2A-B55 activity shown in (C) is the same as shown in Figure 4C, except that we have plotted activity as a function of Cdk1 activity rather than cyclin concentration.

**Figure S6, related to Figure 7.**
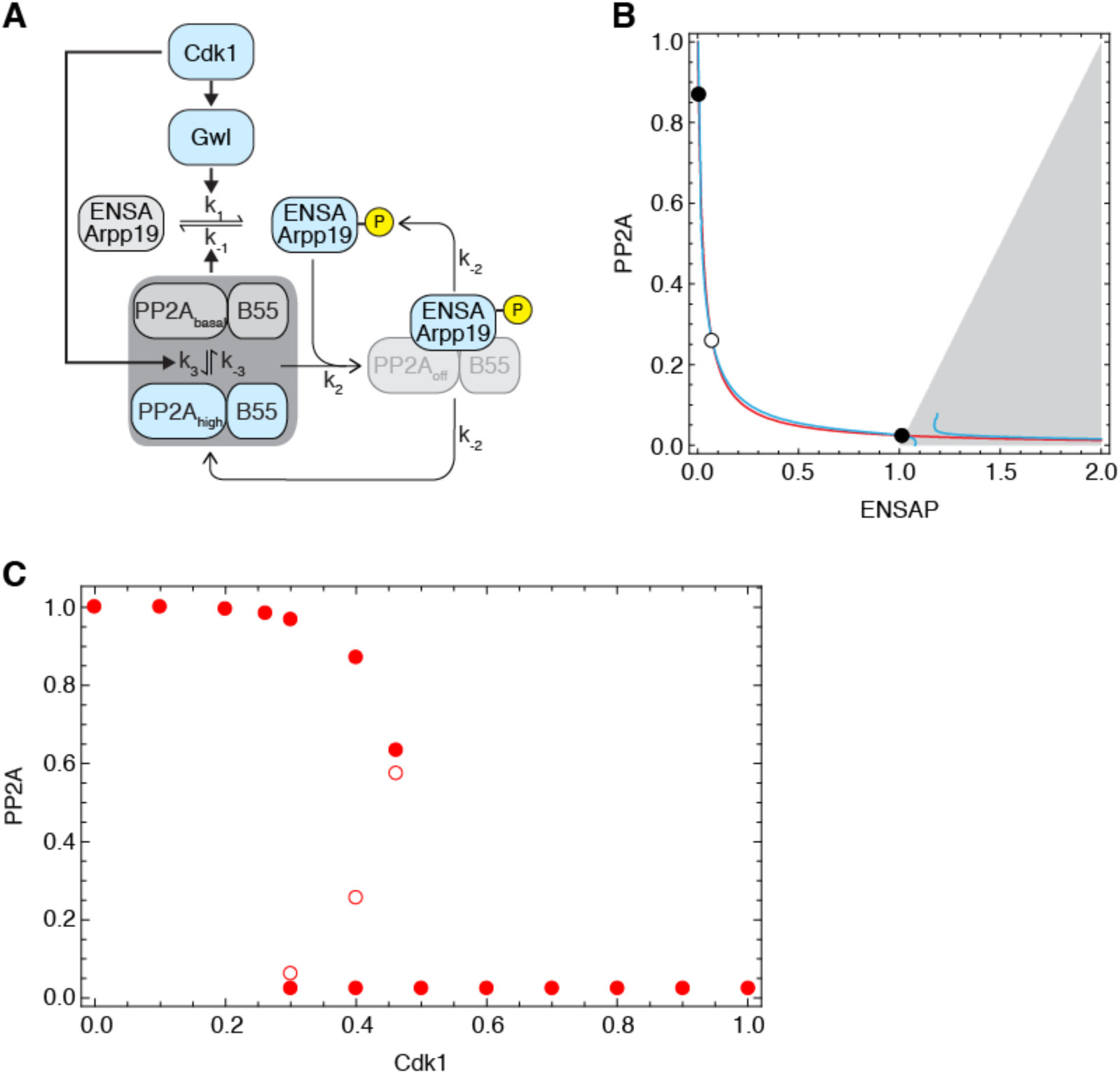
A model for the biphasic, bistable response in the PP2A-B55/ENSA circuit. (A) Schematic depiction of the modelled Cdk1-ENSA/Arpp19-PP2A regulatory circuit. We assume that PP2A-B55 can exist as a basal or high activity form and that Cdk1 promotes the transition into the high activity form. Cdk1 also promotes the phosphorylation of ENSA/Arpp19 via Greatwall, and PP2A dephosphorylates ENSA/Arpp19. Additionally, phosphorylated ENSA can bind to and inhibit both forms of PP2A-B55. (B) Phase plane plot of free PP2A-B55 (both basal and high activity forms) versus phosphorylated ENSA/Arpp19 (ENSAP). The nullclines for PP2A (red) and ENSAP (blue) intersect at three points corresponding to the two stable steady states (black filled circles) and one saddle point (hollow circle). Note that the grey shaded portion of the phase plan is not accessible under the assumption of physiological protein concentrations (> 0 nM). (C) Numerical steady-state solutions (filled circles for stable, hollow circles for unstable) for free PP2A-B55 (basal and high activity) as a function of Cdk1 activity. More than one solution was found for intermediate levels of Cdk1 activity signifying the bistable region.

### Supplemental text

Here we implement a relatively simple model for the regulation of PP2A-B55 by phosphorylated ENSA and Arpp19, the regulation of ENSA and Arpp19 by PP2A-B55, and the regulation of the whole system by active cyclin B-Cdk1. The goal is to account for (1) the bistability of the PP2A-B55/ENSA subsystem; (2) the biphasic dependence of PP2A-B55 activity on cyclin B-Cdk1; and (3) the temporal dynamics of PP2A-B55 regulation, with two peaks of activity per cell cycle.

The basic circuit is shown schematically in Figure S6A. We begin with the core of the system, the double-negative feedback between PP2A-B55 and ENSA/Arpp19. We assume that the system is spatially homogeneous and begin by writing ordinary differential equations (ODEs) describing the time evolution of the key species. For the net production of ENSA/Arpp19 (denoted *ENSA*) we assume that *ENSA* can be produced by the saturable dephosphorylation of *ENSAP*, catalyzed by *PP2A*, and can be consumed by saturable Gwl-mediated phosphorylation. For simplicity we also assume that Gwl is always in steady-state with respect to the activity of cyclin B-Cdk1 (denoted *Cdk1*), and that the steady-state response of Gwl to Cdk1 is ultrasensitive as found experimentally (Figures 4 and S3). Thus,

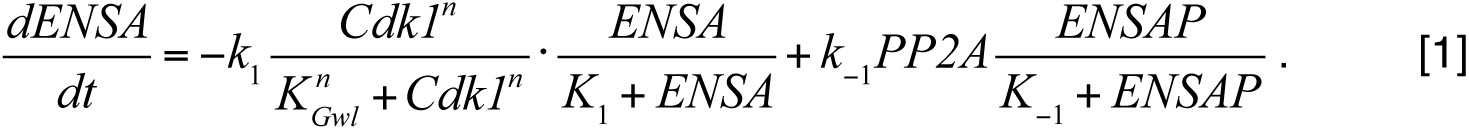

For *ENSAP* we assume that there is one additional way it can be consumed—by binding to and neutralizing *PP2A*—and one additional way it can be produced, by dissociating from *PP2A*.

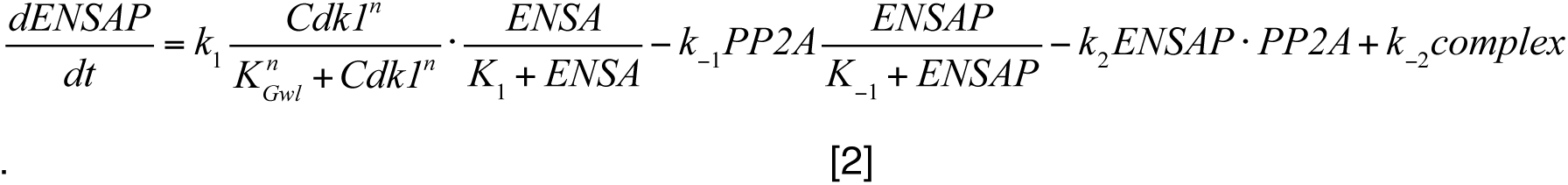

Note that this mechanism supposes that there are two ways ENSAP and PP2A can interact: enzymatically, with a transient interaction resulting in the dephosphorylation of ENSA, and stoichiometrically, where the ENSAP and PP2A complex is essentially inert and the only way for ENSAP to be dephosphorylated or for PP2A to act on any of its substrates is to first dissociate from the complex. This is not the simplest way to envision the interplay between the two proteins (see, for example, Vinod and Novak (Vinod and Novak, 2015)), but it is one of the simplest mechanisms that can give rise to bistability. Further discussion of bistability in double-negative feedback loops with stoichiometric inhibition can be found in Thron (Thron, 1999).

For PP2A there are two issues: the activation of PP2A by Cdk1 (or something downstream of Cdk1) and the binding of PP2A to ENSAP. Here we assume that both the basal-activity form of PP2A and the high-activity form of PP2A can be inhibited equally well by binding to ENSAP. This assumption allows us to write the following ODEs for free PP2A (basal and high activity forms taken together) and for the ENSAP-PP2A complex:

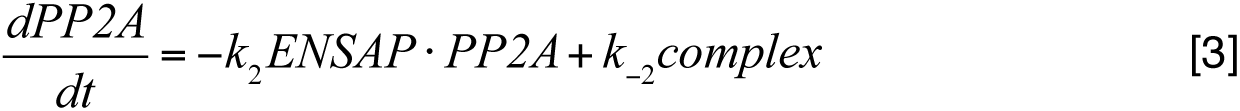

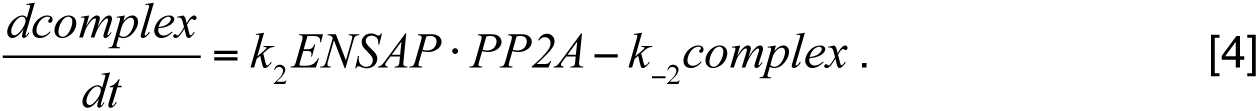

Finally, there are two conservation equations.

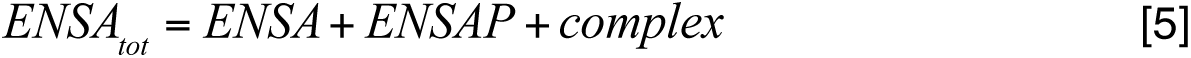

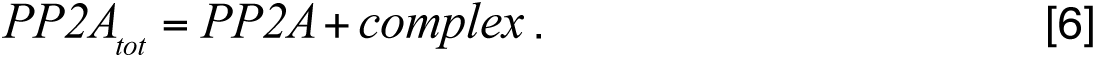

We used these equations to eliminate two variables (*ENSA* and *complex*) from Eqs 2 and 3, which reduces the system to 2 ODEs and 2 time-dependent variables. We plotted the nullclines corresponding to these 2 ODEs and manually searched (using the `Manipulate` command in Mathematica) for parameters that yielded three intersection points between the nullclines and verified that they corresponded to two stable steady states and one saddle point. We also adjusted the nullclines so that the concentration of free PP2A would be substantially different (∼10-fold different) between the two stable steady states.

The phase plane plot of the nullclines and steady states is shown in Figure S6B. Note that despite the fact that the two null clines run very close to each other, the resulting bistable response is surprisingly robust, with a realistic degree of hysteresis (Figure S6C). Note as well that part of the phase plane (the shaded portion) is not accessible if all of the concentrations are required to be positive quantities.

We then calculated the positions of the steady states as a function of the Cdk1 concentration. Mathematica was able to derive closed-form expressions for *ENSAP* and *PP2A* as functions of the parameters and Cdk1, but it was so large and unwieldy that we elected instead to solve for the steady states numerically for a series of assumed values of Cdk1. The resulting curve (shown for a course grid of Cdk1 values) is shown in Figure S6C.

To convert these concentrations of free PP2A to activities, we assumed that the conversion of PP2A between its basal and active forms was mediated by phosphorylation, with Cdk1 being the activating kinase and some constitutive phosphatase carrying out the inactivating dephosphorylation. This yields the following ODE:

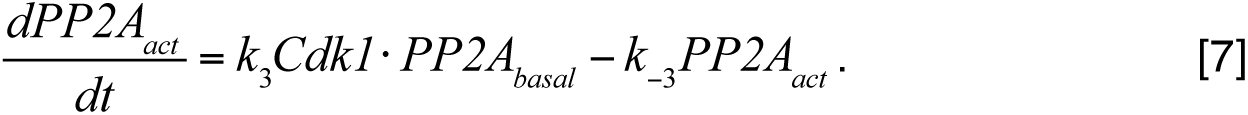

Using the conservation relationship *PP2A* = *PP2A_basal_*+ *PP2A_act_* and solving for the two forms of PP2A at steady state yields:

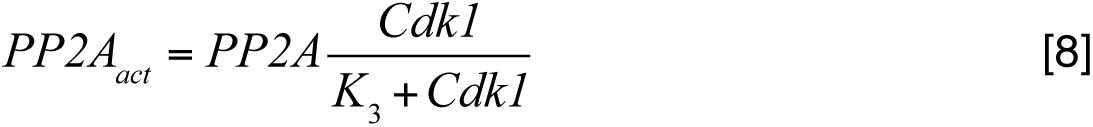

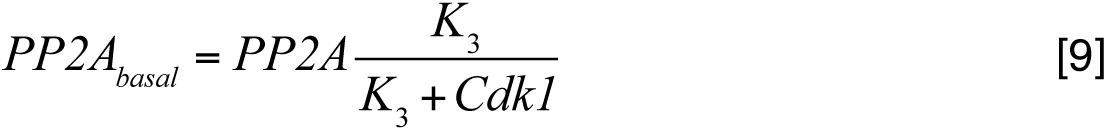

where *K_3_* = *k*_−_*_3_ / k_3_*. The total PP2A activity is then taken to be:

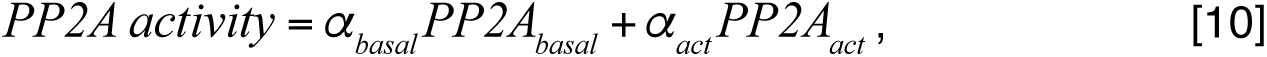

where the weighting factors (*α_basal_* and *α_act_*) define the relative activities of the two forms of PP2A. This yielded the activity vs. Cdk1 curve shown in Figure 7G.

To calculate substrate phosphorylation at steady state, we assumed that the process was described by mass action kinetics:

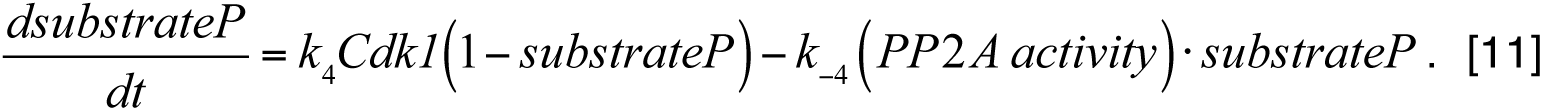

At steady state this yields a Michaelian (hyperbolic response). This was used for the phosphorylation vs. Cdk1 curve shown in Figure 7H.

Finally, to calculate the time course of PP2A activation and inactivation and substrate phosphorylation, we assumed that Cdk1 activity initially rose gradually, then rose more steeply, and then fell exponentially during mitotic exit. We then solved the 2 ODE system (Eqs 2 and 3) numerically and compared the calculated time courses for PP2A activity and substrate phosphorylation to the experimental data in Figure 7E. We found that to obtain realistic-looking time courses, we needed to assume that the processes described above were rapid compared to the changes in Cdk1. We multiplied all of the rate constants in the rate equations by an appropriate scaling factor; the final parameter values are listed in Supplementary Table 1. This yielded the time course simulations shown in Figure 7F.

In summary, this model accounts for the biphasic nature of PP2A regulation, the bistability of PP2A inactivation, and the qualitative features of the time course experiments.

### Supplemental table

**Table S1 related to Figure S6.** Supplemental text. Model parameters.

| Parameter | Meaning | Value |
| --- | --- | --- |
| $k_1$ | Rate constant for the phosphorylation of ENSA | 15 |
| $K_1$ | Michaelis constant for the phosphorylation of ENSA | 0.1 |
| $k_{-1}$ | Rate constant for the dephosphorylation of ENSAP | 1.5 |
| $K_{-1}$ | Michaelis constant for the dephosphorylation of ENSAP | 0.01 |
| $k_2$ | Rate constant for the binding of ENSAP to PP2A | 600 |
| $k_{-2}$ | Rate constant for the dissociation of ENSAP from PP2 | 15 |
| $n$ | Hill exponent for Gwl activation by Cdk1 | 5 |
| $K_3$ | EC50 for PP2A activation by Cdk1 | 10 |
| $\alpha_{basal}$ | Weighting factor for basal PP2A | 0.3 |
| $\alpha_{act}$ | Weighting factor for active PP2A | 15 |
| $k_4$ | Rate constant for substrate phosphorylation | 0.4 |
| $k_{-4}$ | Rate constant for substrate dephosphorylation | 2 |
| $ENSA_{tot}$ | Total concentration of ENSA | 2 |
| $PP2A_{tot}$ | Total concentration of PP2A | 1 |

## References

1. Morgan, D.O. (2007). The Cell Cycle: Principles of Control., (New Science Press).

2. Swaffer, M.P., Jones, A.W., Flynn, H.R., Snijders, A.P., and Nurse, P. (2018). Quantitative Phosphoproteomics Reveals the Signaling Dynamics of Cell-Cycle Kinases in the Fission Yeast Schizosaccharomyces pombe. Cell Rep 24, 503–514.

3. Dephoure, N., Zhou, C., Villen, J., Beausoleil, S.A., Bakalarski, C.E., Elledge, S.J., and Gygi, S.P. (2008). A quantitative atlas of mitotic phosphorylation. Proc Natl Acad Sci U S A 105, 10762–10767.

4. Ubersax, J.A., Woodbury, E.L., Quang, P.N., Paraz, M., Blethrow, J.D., Shah, K., Shokat, K.M., and Morgan, D.O. (2003). Targets of the cyclin-dependent kinase Cdk1. Nature 425, 859–864.

5. Healy, A.M., Zolnierowicz, S., Stapleton, A.E., Goebl, M., DePaoli-Roach, A.A., and Pringle, J.R. (1991). CDC55, a Saccharomyces cerevisiae gene involved in cellular morphogenesis: identification, characterization, and homology to the B subunit of mammalian type 2A protein phosphatase. Molecular and cellular biology 11, 5767–5780.

6. Mochida, S., Ikeo, S., Gannon, J., and Hunt, T. (2009). Regulated activity of PP2A-B55 delta is crucial for controlling entry into and exit from mitosis in Xenopus egg extracts. The EMBO journal 28, 2777–2785.

7. Schmitz, M.H., Held, M., Janssens, V., Hutchins, J.R., Hudecz, O., Ivanova, E., Goris, J., Trinkle-Mulcahy, L., Lamond, A.I., Poser, I., et al. (2010). Live-cell imaging RNAi screen identifies PP2A-B55alpha and importin-beta1 as key mitotic exit regulators in human cells. Nature cell biology 12, 886–893.

8. Wu, J.Q., Guo, J.Y., Tang, W., Yang, C.S., Freel, C.D., Chen, C., Nairn, A.C., and Kornbluth, S. (2009). PP1-mediated dephosphorylation of phosphoproteins at mitotic exit is controlled by inhibitor-1 and PP1 phosphorylation. Nature cell biology 11, 644–651.

9. Akopyan, K., Silva Cascales, H., Hukasova, E., Saurin, A.T., Mullers, E., Jaiswal, H., Hollman, D.A., Kops, G.J., Medema, R.H., and Lindqvist, A. (2014). Assessing kinetics from fixed cells reveals activation of the mitotic entry network at the S/G2 transition. Mol Cell 53, 843–853.

10. Murray, A.W., and Kirschner, M.W. (1989). Cyclin synthesis drives the early embryonic cell cycle. Nature 339, 275–280.

11. Solomon, M.J., Glotzer, M., Lee, T.H., Philippe, M., and Kirschner, M.W. (1990). Cyclin activation of p34cdc2. Cell 63, 1013–1024.

12. Sunkara, P.S., Wright, D.A., and Rao, P.N. (1979). Mitotic factors from mammalian cells induce germinal vesicle breakdown and chromosome condensation in amphibian oocytes. Proc Natl Acad Sci U S A 76, 2799–2802.

13. Novak, B., and Tyson, J.J. (1993). Modeling the Cell-Division Cycle - M-Phase Trigger, Oscillations, and Size Control. J Theor Biol 165, 101–134.

14. Goldbeter, A. (1993). Modeling the mitotic oscillator driving the cell division cycle. Comments on Theoretical Biology 3, 75–107.

15. Novak, B., and Tyson, J.J. (1993). Numerical analysis of a comprehensive model of M-phase control in Xenopus oocyte extracts and intact embryos. J Cell Sci 106 ( *Pt 4*), 1153–1168.

16. Gould, K.L., and Nurse, P. (1989). Tyrosine phosphorylation of the fission yeast cdc2+ protein kinase regulates entry into mitosis. Nature 342, 39–45.

17. Krek, W., and Nigg, E.A. (1991). Differential phosphorylation of vertebrate p34cdc2 kinase at the G1/S and G2/M transitions of the cell cycle: identification of major phosphorylation sites. The EMBO journal 10, 305–316.

18. McGowan, C.H., and Russell, P. (1993). Human Wee1 kinase inhibits cell division by phosphorylating p34cdc2 exclusively on Tyr15. The EMBO journal 12, 75–85.

19. Mueller, P.R., Coleman, T.R., Kumagai, A., and Dunphy, W.G. (1995). Myt1: a membrane-associated inhibitory kinase that phosphorylates Cdc2 on both threonine-14 and tyrosine-15. Science 270, 86–90.

20. Parker, L.L., and Piwnica-Worms, H. (1992). Inactivation of the p34cdc2-cyclin B complex by the human WEE1 tyrosine kinase. Science 257, 1955–1957.

21. Millar, J.B., McGowan, C.H., Lenaers, G., Jones, R., and Russell, P. (1991). p80cdc25 mitotic inducer is the tyrosine phosphatase that activates p34cdc2 kinase in fission yeast. The EMBO journal 10, 4301–4309.

22. Strausfeld, U., Labbe, J.C., Fesquet, D., Cavadore, J.C., Picard, A., Sadhu, K., Russell, P., and Doree, M. (1991). Dephosphorylation and activation of a p34cdc2/cyclin B complex in vitro by human CDC25 protein. Nature 351, 242–245.

23. Alfieri, C., Zhang, S., and Barford, D. (2017). Visualizing the complex functions and mechanisms of the anaphase promoting complex/cyclosome (APC/C). Open Biol 7.

24. Peters, J.M. (2006). The anaphase promoting complex/cyclosome: a machine designed to destroy. Nat Rev Mol Cell Biol 7, 644–656.

25. Glotzer, M., Murray, A.W., and Kirschner, M.W. (1991). Cyclin is degraded by the ubiquitin pathway. Nature 349, 132–138.

26. Hershko, A., Ganoth, D., Pehrson, J., Palazzo, R.E., and Cohen, L.H. (1991). Methylated ubiquitin inhibits cyclin degradation in clam embryo extracts. J Biol Chem 266, 16376–16379.

27. King, R.W., Peters, J.M., Tugendreich, S., Rolfe, M., Hieter, P., and Kirschner, M.W. (1995). A 20S complex containing CDC27 and CDC16 catalyzes the mitosis-specific conjugation of ubiquitin to cyclin B. Cell 81, 279–288.

28. Yamano, H., Ishii, K., and Yanagida, M. (1994). Phosphorylation of dis2 protein phosphatase at the C-terminal cdc2 consensus and its potential role in cell cycle regulation. The EMBO journal 13, 5310–5318.

29. Mochida, S., Maslen, S.L., Skehel, M., and Hunt, T. (2010). Greatwall phosphorylates an inhibitor of protein phosphatase 2A that is essential for mitosis. Science 330, 1670–1673.

30. Vigneron, S., Brioudes, E., Burgess, A., Labbe, J.C., Lorca, T., and Castro, A. (2009). Greatwall maintains mitosis through regulation of PP2A. The EMBO journal 28, 2786–2793.

31. Nasa, I., Cressey, L.E., Kruse, T., Hertz, E.P.T., Gui, J., Graves, L.M., Nilsson, J., and Kettenbach, A.N. (2020). Quantitative kinase and phosphatase profiling reveal that CDK1 phosphorylates PP2Ac to promote mitotic entry. Sci Signal 13.

32. Vinod, P.K., and Novak, B. (2015). Model scenarios for switch-like mitotic transitions. FEBS Lett 589, 667–671.

33. Mochida, S., Rata, S., Hino, H., Nagai, T., and Novak, B. (2016). Two Bistable Switches Govern M Phase Entry. Curr Biol 26, 3361–3367.

34. Rata, S., Suarez Peredo Rodriguez, M.F., Joseph, S., Peter, N., Echegaray Iturra, F., Yang, F., Madzvamuse, A., Ruppert, J.G., Samejima, K., Platani, M., et al. (2018). Two Interlinked Bistable Switches Govern Mitotic Control in Mammalian Cells. Curr Biol 28, 3824–3832 e3826.

35. Pomerening, J.R., Sontag, E.D., and Ferrell, J.E., Jr. (2003). Building a cell cycle oscillator: hysteresis and bistability in the activation of Cdc2. Nature cell biology 5, 346–351.

36. Sha, W., Moore, J., Chen, K., Lassaletta, A.D., Yi, C.S., Tyson, J.J., and Sible, J.C. (2003). Hysteresis drives cell-cycle transitions in Xenopus laevis egg extracts. Proc Natl Acad Sci U S A 100, 975–980.

37. Ferrell, J.E., Jr., Wu, M., Gerhart, J.C., and Martin, G.S. (1991). Cell cycle tyrosine phosphorylation of p34cdc2 and a microtubule-associated protein kinase homolog in Xenopus oocytes and eggs. Molecular and cellular biology 11, 1965–1971.

38. Hartley, R.S., Rempel, R.E., and Maller, J.L. (1996). In vivo regulation of the early embryonic cell cycle in Xenopus. Developmental biology 173, 408–419.

39. Tsai, T.Y., Theriot, J.A., and Ferrell, J.E., Jr. (2014). Changes in oscillatory dynamics in the cell cycle of early Xenopus laevis embryos. PLoS biology 12, e1001788.

40. Chang, J.B., and Ferrell, J.E., Jr. (2013). Mitotic trigger waves and the spatial coordination of the Xenopus cell cycle. Nature 500, 603–607.

41. Coudreuse, D., and Nurse, P. (2010). Driving the cell cycle with a minimal CDK control network. Nature 468, 1074–1079.

42. Navarro, F.J., and Nurse, P. (2012). A systematic screen reveals new elements acting at the G2/M cell cycle control. Genome Biol 13, R36.

43. Kim, S.Y., and Ferrell, J.E., Jr. (2007). Substrate competition as a source of ultrasensitivity in the inactivation of Wee1. Cell 128, 1133–1145.

44. Trunnell, N.B., Poon, A.C., Kim, S.Y., and Ferrell, J.E., Jr. (2011). Ultrasensitivity in the Regulation of Cdc25C by Cdk1. Mol Cell 41, 263–274.

45. Yang, Q., and Ferrell, J.E., Jr. (2013). The Cdk1-APC/C cell cycle oscillator circuit functions as a time-delayed, ultrasensitive switch. Nature cell biology 15, 519–525.

46. Wang, Y., Li, J., Booher, R.N., Kraker, A., Lawrence, T., Leopold, W.R., and Sun, Y. (2001). Radiosensitization of p53 mutant cells by PD0166285, a novel G(2) checkpoint abrogator. Cancer Res 61, 8211–8217.

47. Shou, W., and Dunphy, W.G. (1996). Cell cycle control by Xenopus p28Kix1, a developmentally regulated inhibitor of cyclin-dependent kinases. Molecular biology of the cell 7, 457–469.

48. Georgi, A.B., Stukenberg, P.T., and Kirschner, M.W. (2002). Timing of events in mitosis. Curr Biol 12, 105–114.

49. Fujimitsu, K., Grimaldi, M., and Yamano, H. (2016). Cyclin-dependent kinase 1-dependent activation of APC/C ubiquitin ligase. Science 352, 1121–1124.

50. Yamaguchi, M., Yu, S., Qiao, R., Weissmann, F., Miller, D.J., VanderLinden, R., Brown, N.G., Frye, J.J., Peters, J.M., and Schulman, B.A. (2015). Structure of an APC3-APC16 complex: insights into assembly of the anaphase-promoting complex/cyclosome. J Mol Biol 427, 1748-1764.

51. Zhang, S., Chang, L., Alfieri, C., Zhang, Z., Yang, J., Maslen, S., Skehel, M., and Barford, D. (2016). Molecular mechanism of APC/C activation by mitotic phosphorylation. Nature 533, 260–264.

52. Tischer, T., Hormanseder, E., and Mayer, T.U. (2012). The APC/C inhibitor XErp1/Emi2 is essential for Xenopus early embryonic divisions. Science 338, 520–524.

53. Vinod, P.K., Zhou, X., Zhang, T., Mayer, T.U., and Novak, B. (2013). The role of APC/C inhibitor Emi2/XErp1 in oscillatory dynamics of early embryonic cell cycles. Biophysical chemistry 177-178, 1–6.

54. Dohadwala, M., da Cruz e Silva, E.F., Hall, F.L., Williams, R.T., Carbonaro-Hall, D.A., Nairn, A.C., Greengard, P., and Berndt, N. (1994). Phosphorylation and inactivation of protein phosphatase 1 by cyclin-dependent kinases. Proc Natl Acad Sci U S A 91, 6408–6412.

55. Kwon, Y.G., Lee, S.Y., Choi, Y., Greengard, P., and Nairn, A.C. (1997). Cell cycle-dependent phosphorylation of mammalian protein phosphatase 1 by cdc2 kinase. Proc Natl Acad Sci U S A 94, 2168–2173.

56. Yu, J., Zhao, Y., Li, Z., Galas, S., and Goldberg, M.L. (2006). Greatwall kinase participates in the Cdc2 autoregulatory loop in Xenopus egg extracts. Mol Cell 22, 83–91.

57. Zhao, Y., Haccard, O., Wang, R., Yu, J., Kuang, J., Jessus, C., and Goldberg, M.L. (2008). Roles of Greatwall kinase in the regulation of cdc25 phosphatase. Molecular biology of the cell 19, 1317–1327.

58. Blake-Hodek, K.A., Williams, B.C., Zhao, Y., Castilho, P.V., Chen, W., Mao, Y., Yamamoto, T.M., and Goldberg, M.L. (2012). Determinants for activation of the atypical AGC kinase Greatwall during M phase entry. Molecular and cellular biology 32, 1337–1353.

59. Gharbi-Ayachi, A., Labbe, J.C., Burgess, A., Vigneron, S., Strub, J.M., Brioudes, E., Van-Dorsselaer, A., Castro, A., and Lorca, T. (2010). The substrate of Greatwall kinase, Arpp19, controls mitosis by inhibiting protein phosphatase 2A. Science 330, 1673–1677.

60. Mochida, S., and Hunt, T. (2012). Protein phosphatases and their regulation in the control of mitosis. EMBO reports 13, 197–203.

61. Williams, B.C., Filter, J.J., Blake-Hodek, K.A., Wadzinski, B.E., Fuda, N.J., Shalloway, D., and Goldberg, M.L. (2014). Greatwall-phosphorylated Endosulfine is both an inhibitor and a substrate of PP2A-B55 heterotrimers. Elife 3, e01695.

62. Hegarat, N., Vesely, C., Vinod, P.K., Ocasio, C., Peter, N., Gannon, J., Oliver, A.W., Novak, B., and Hochegger, H. (2014). PP2A/B55 and Fcp1 regulate Greatwall and Ensa dephosphorylation during mitotic exit. PLoS Genet 10, e1004004.

63. Heim, A., Konietzny, A., and Mayer, T.U. (2015). Protein phosphatase 1 is essential for Greatwall inactivation at mitotic exit. EMBO reports 16, 1501-1510.

64. Ma, S., Vigneron, S., Robert, P., Strub, J.M., Cianferani, S., Castro, A., and Lorca, T. (2016). Greatwall dephosphorylation and inactivation upon mitotic exit is triggered by PP1. J Cell Sci 129, 1329-1339.

65. Rogers, S., Fey, D., McCloy, R.A., Parker, B.L., Mitchell, N.J., Payne, R.J., Daly, R.J., James, D.E., Caldon, C.E., Watkins, D.N., et al. (2016). PP1 initiates the dephosphorylation of MASTL, triggering mitotic exit and bistability in human cells. J Cell Sci 129, 1340–1354.

66. Mochida, S., and Hunt, T. (2007). Calcineurin is required to release Xenopus egg extracts from meiotic M phase. Nature 449, 336–340.

67. Bialojan, C., and Takai, A. (1988). Inhibitory effect of a marine-sponge toxin, okadaic acid, on protein phosphatases. Specificity and kinetics. Biochem J 256, 283–290.

68. Mochida, S. (2014). Regulation of alpha-endosulfine, an inhibitor of protein phosphatase 2A, by multisite phosphorylation. FEBS J 281, 1159–1169.

69. Wuhr, M., Freeman, R.M., Jr., Presler, M., Horb, M.E., Peshkin, L., Gygi, S., and Kirschner, M.W. (2014). Deep proteomics of the Xenopus laevis egg using an mRNA-derived reference database. Curr Biol 24, 1467–1475.

70. Hopkins, M., Tyson, J.J., and Novak, B. (2017). Cell-cycle transitions: a common role for stoichiometric inhibitors. Molecular biology of the cell 28, 3437–3446.

71. Thron, C.D. (1999). Mathematical analysis of binary activation of a cell cycle kinase which down-regulates its own inhibitor. Biophysical chemistry 79, 95–106.

72. Mangan, S., and Alon, U. (2003). Structure and function of the feed-forward loop network motif. Proc Natl Acad Sci U S A 100, 11980–11985.

73. Tsai, T.Y., Choi, Y.S., Ma, W., Pomerening, J.R., Tang, C., and Ferrell, J.E., Jr. (2008). Robust, tunable biological oscillations from interlinked positive and negative feedback loops. Science 321, 126–129.

74. Gelens, L., and Saurin, A.T. (2018). Exploring the Function of Dynamic Phosphorylation-Dephosphorylation Cycles. Dev Cell 44, 659–663.

75. Rodenfels, J., Neugebauer, K.M., and Howard, J. (2019). Heat Oscillations Driven by the Embryonic Cell Cycle Reveal the Energetic Costs of Signaling. Dev Cell.

76. Pomerening, J.R., Kim, S.Y., and Ferrell, J.E., Jr. (2005). Systems-level dissection of the cell-cycle oscillator: bypassing positive feedback produces damped oscillations. Cell 122, 565–578.

77. Murray, A.W., Solomon, M.J., and Kirschner, M.W. (1989). The role of cyclin synthesis and degradation in the control of maturation promoting factor activity. Nature 339, 280–286.

78. Kumagai, A., and Dunphy, W.G. (1992). Regulation of the cdc25 protein during the cell cycle in Xenopus extracts. Cell 70, 139–151.

79. Draetta, G., and Beach, D. (1988). Activation of cdc2 protein kinase during mitosis in human cells: cell cycle-dependent phosphorylation and subunit rearrangement. Cell 54, 17–26.

80. Swaffer, M.P., Jones, A.W., Flynn, H.R., Snijders, A.P., and Nurse, P. (2016). CDK Substrate Phosphorylation and Ordering the Cell Cycle. Cell 167, 1750–1761 e1716.

81. Longin, S., Zwaenepoel, K., Louis, J.V., Dilworth, S., Goris, J., and Janssens, V. (2007). Selection of protein phosphatase 2A regulatory subunits is mediated by the C terminus of the catalytic Subunit. J Biol Chem 282, 26971–26980.

82. Vigneron, S., Sundermann, L., Labbe, J.C., Pintard, L., Radulescu, O., Castro, A., and Lorca, T. (2018). Cyclin A-cdk1-Dependent Phosphorylation of Bora Is the Triggering Factor Promoting Mitotic Entry. Dev Cell 45, 637–650 e637.

83. Murray, A. (1991). Cell Cycle Extracts, Volume 36.

84. Vollmer, B., Schooley, A., Sachdev, R., Eisenhardt, N., Schneider, A.M., Sieverding, C., Madlung, J., Gerken, U., Macek, B., and Antonin, W. (2012). Dimerization and direct membrane interaction of Nup53 contribute to nuclear pore complex assembly. The EMBO journal 31, 4072–4084.

85. Kim, S.Y., Song, E.J., Lee, K.J., and Ferrell, J.E., Jr. (2005). Multisite M-phase phosphorylation of Xenopus Wee1A. Molecular and cellular biology 25, 10580–10590.

